# Slowdown of microtubule retrograde flow enables axon and dendrite development and maintenance

**DOI:** 10.64898/2025.12.18.694605

**Authors:** Max Schelski, Thorben Pietralla, Cedric Günter, Sina Stern, Nenad Pavin, Frank Bradke

## Abstract

Distinct microtubule arrays form in axons and dendrites defining their functions in neurons. These arrays are thought to develop through separate processes. Here we challenge this view by showing that axons and dendrites develop by a unifying process: slowdown of microtubule retrograde flow (MT-RF), a recently discovered mechanism of microtubule dynamics. By integrating quantitative data of microtubule dynamics and distributions into a newly developed biophysical model of microtubules across developmental stages, we uncover that MT-RF interacts with microtubule stability and nucleation. Without MT-RF slowdown, this interaction suppresses development and maintenance of axonal and dendritic microtubule arrays. In axons, MT-RF slowdown enables microtubules to reach the tip, even at long distances, supporting axon growth. In dendrites, late MT-RF slowdown facilitates an efficient increase in stable microtubules. Thus, our data-integrated model reveals MT-RF slowdown as central for the development of distinct neuronal compartments, providing a unifying mechanism for microtubule-related physiology.

---

Axons and dendrites differ in their morphology and molecular composition; this polarization enables neurons to function^1,2^. Differences between axons and dendrites are thought to be established through separate developmental processes^2^. Developing neurites, axons, and dendrites differ in their microtubule (MT) array^3,4^. In developing neurites, the majority of MTs orient with their growing plus-end towards the neurite tip (plus-end-out), and a smaller fraction of MTs orients towards the soma (plus-end-in)^5^. In axons, MTs form a uniformly oriented array with plus-end-out MTs^5–7^, while the MT array in dendrites consists of MTs with both orientations in equal parts^5,6^. These MT differences direct polarized transport into axons and dendrites^8,9^. This has led to a large body of research addressing which separate mechanisms in either axons or dendrites enable such a distinct organization^3,8–11^.

Here, we asked instead whether the axonal and dendritic MT arrays could be created through one central MT regulatory mechanism. We hypothesized that such a central mechanism could provide specific and distinct MT arrays through interactions with a minimal number of MT modulations. To identify such a putative central mechanism, we focused on common characteristics of neuronal MT arrays. First, developing neurites, axons, and dendrites all rely on MTs to grow^12–14^. Second, these neuronal compartments depend on stable MTs. In axons, the amount of stable MTs increases compared to developing neurites, and this increase is sufficient to induce axon development^15^. In dendrites, stable and unstable MTs are differently oriented^9^. Stable MTs are predominantly plus-end-in oriented thereby enabling cargo-sorting by directing axon-specific cargo out of dendrites^9^. Third, we and others recently found that, in developing neurites, unexpectedly, the entire MT array moves retrogradely towards the soma^16,17^. This MT retrograde flow (MT-RF) slows down in the axon and later also in dendrites^16,17^. Could these similarities indicate a common mechanism driving neuronal polarity?

By building a biophysical model integrating experimental data across developmental stages, we identified slowdown of MT-RF as the unifying mechanism for the development and maintenance of axons, and dendrites. MT-RF slowdown, together with only three modulations of the MT array – stability and two types of nucleation – explains the formation of the stable MT distribution in axons and dendrites, which gives rise to polarized neuronal morphology.

## Results

### Stable and unstable MTs have different distributions in developing neurites

MTs across the length of axons and dendrites have several functions. For example, distal MTs at the axon tip drive axon growth^12,15,18–20^, while the distribution of MTs across the dendrite directs transport^10^. To investigate the regulation of the MT array during development, we characterized the distribution of stable and unstable MTs in developing neurites after one day in culture. We first assessed the fraction of stable MT mass by measuring the decay of MT mass after photoconversion in the proximal neurite, when MTs moved retrogradely to the cell body, referred to as MT-RF^16,17^ (Fig. 1a-c). The decay curve of photoconverted MT mass was fitted well by a dual-exponential function but not a mono-exponential function (Fig. 1d). This indicated the presence of two populations of MTs with distinct decay constants. We observed a fast-decaying component that corresponds to unstable MTs and a slow-decaying component that corresponds to stable MTs (Fig. 1b,d). In 84% of neurites, a part of MT mass decayed slowly (stable MTs). Overall, 40 ± 4% of MT mass decayed slowly with the median half-life of 9.1 min while 60 ± 4% of MT mass decayed fast with a median half-life of 1.0 min (unstable MTs; mean ± SEM; Fig. 1e-f).

**Fig. 1.**
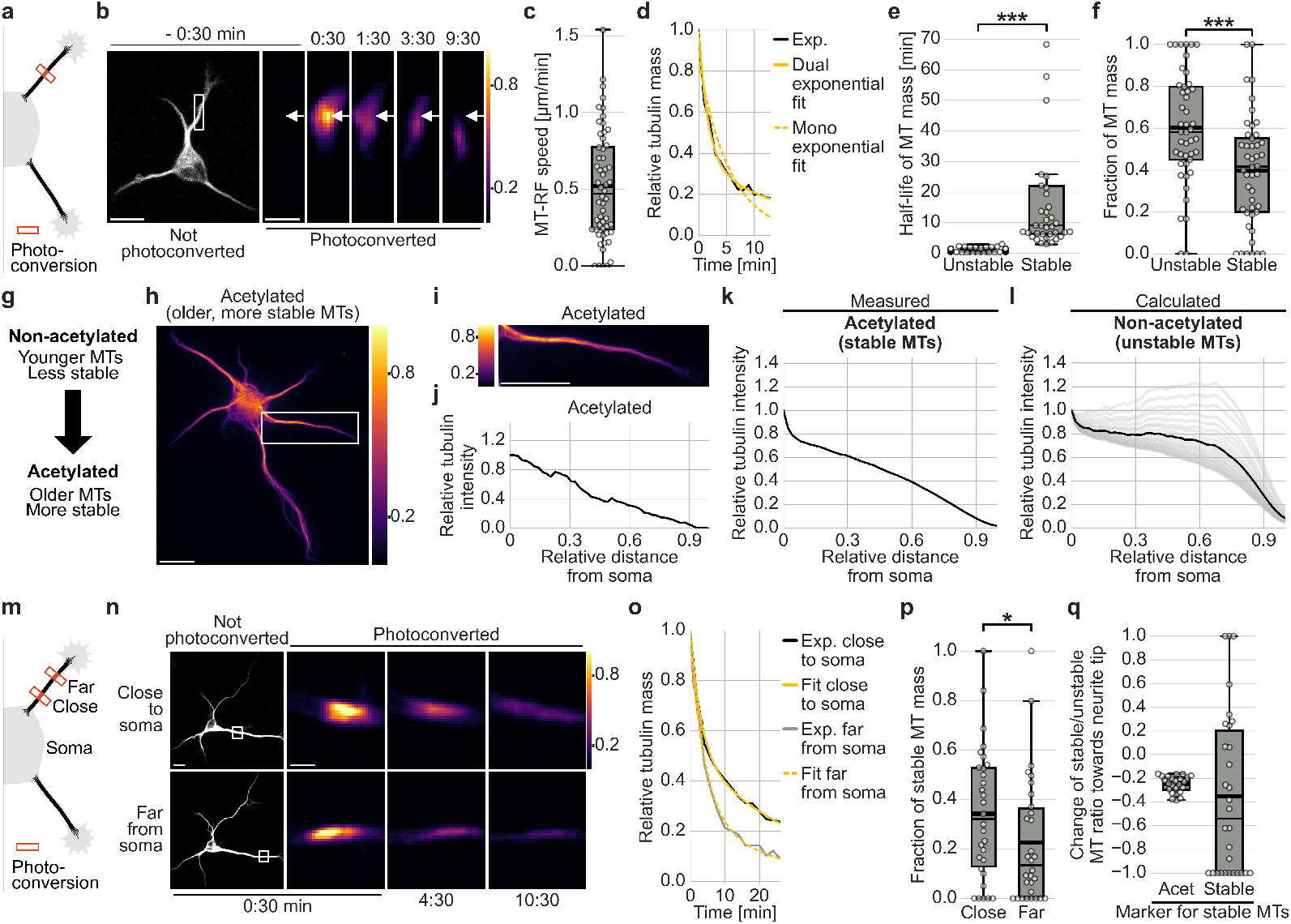
Stable and unstable MTs are differently distributed in developing neurites. **a-f**, The tubulin TUBB2a was fused to the photoconvertible fluorophore mEos3.2 and expressed in neurons in which a small MT patch was photoconverted after one day in culture. (**a**) Illustration of photoconversion experiment for **b-f**. (**b**) Representative images of the not photoconverted channel and the photoconverted channel from before to after photoconversion for **c-f**. The white arrow marks the position of the photoconverted MT array at the start. Time is in min:seconds. Scale bars, 10 µm for not zoomed and 2 µm for zoomed images. (**c**) MT-RF was measured as the distance that the photoconverted MT patch moved per time (n = 48 neurons, N = 4 experiments). (**d-f**) The photoconverted fluorescent intensity in the entire neurite was summed at each timepoint. (**d**) The resulting decay curve (Exp.) was then fitted to either a mono- or dual-exponential function, which indicates one population or two populations of decaying MTs. (**e,f**) (**e**) Half-life and (**f**) fraction of the photoconverted MT mass of the fast-decaying component (unstable MTs) and the slow decaying component (stable MTs) of the dual exponential fit were plotted. For stable MT half-life in **e** three datapoints with a half-life above 10,000 min are out of the plotting range. (**e**: n = 38 neurons for stable and n = 43 neurons for unstable MTs, N = 4, p = 5.5 * 10^-15^; **f**: n = 45 neurons for both groups, N = 4, p=0.0003; Mann-Whitney U test). **g-l**, Neurons were stained after one day in culture for tyrosination as a marker for younger and detyrosination as well as acetylation as a marker for older MTs. Intensities were quantified for each point along the neurite as the sum of all pixel intensities across the width of the neurite. Tubulin intensity was normalized to the value closest to the soma. (**g**) Illustration of MT acetylation indicating older, more stable MTs. (**h-j**) Representative image (**h**) of the entire neuron and (**i**) the region marked with the white rectangle enlarged, and (**j**) the intensity curve along the neurite from **i**, stained for acetylated tubulin. Scale bars, 10 μm. (**k**) Distributions of tubulin modifications were averaged for all neurites (n = 505 neurites from 143 neurons, N = 3 experiments). (**l**) Based on average distributions of tyrosinated, detyrosinated and acetylated tubulin as well as assumptions about the fraction of co-occurrence of MT modifications, several possible distributions of non-acetylated tubulin were calculated (grey). The average of the possible distributions for non-acetylated tubulin is plotted in black (Methods section “Calculating non-acetylated tubulin distribution”). **m-q**, In neurons expressing TUBB2a fused to mEos3.2, a small area of MTs was photoconverted close to or far from the soma, at an average 30% and 68% neurite length towards the neurite tip, respectively. The photoconverted intensity was summed for each timepoint and the decay curve fitted to a dual exponential function. (**m**) Illustration of photoconversion experiment. (**n**) Images of photoconverted tubulin of a representative neuron for **p**. Time is in min:seconds. Scale bars, 10 µm for not zoomed and 2 µm for zoomed images. (**o**) Decay curves of the representative neuron in **n**, one photoconverted close to and one far from the soma (Exp.) with the corresponding dual exponential fits. (**p**) The fraction of stable MT mass close to and far from the soma (n = 30 neurons, N = 3 experiments; p = 0.029, Mann-Whitney U test). (**q**) The normalized change of the ratio of stable to unstable MTs from close to the soma to far from the soma was calculated and normalized by the sum of the ratios at both positions. Acetylated and non-acetylated tubulin were used as markers for stable and unstable MTs, respectively (Acet). This was compared to the value obtained from the data in **p** as a good marker for stable MTs (Stable). Intensity in **b, h, i** and **n** is color-coded from purple (low) to yellow (high).The thick line in boxplots indicates the mean, the thin line the median.

To measure the distribution of these stable and unstable MTs in neurons, we stained primary hippocampal neurons for acetylated and detyrosinated tubulin as markers for long-lived and thus stable MTs and tyrosinated tubulin as a marker for short-lived and thus unstable MTs^15,21–23^ (Fig. 1g-k; Extended Data Fig. 1a-j). Tyrosinated MTs decreased towards the neurite tip (Extended Data Fig. 1b, e-g). Interestingly, despite both being established markers for stable MTs in neurons^11,15,24,25^, the distributions of acetylated and detyrosinated MTs differed. This difference may occur because detyrosination is slower than acetylation^26^, raising the possibility that they may mark a different subset of MTs. Acetylated MTs decreased towards the neurite tip while detyrosinated MTs remained relatively constant in the first half of the neurite and then decreased (Extended Data Fig. 1b, e-j).

To understand which marker represents stable MTs in neurons, we analyzed the ratio of stable to unstable MTs across the neurite separately for detyrosinated and acetylated tubulin as markers for stable MTs. For acetylated tubulin as a marker for stable MTs, the corresponding marker for unstable MTs would be non-acetylated tubulin. However, since it is not possible to stain for non-acetylated tubulin, we calculated this distribution based on the other tubulin modifications. To this end, we assumed different fractions of MTs that are both acetylated and detyrosinated or acetylated and tyrosinated (Methods section “Calculating non-acetylated tubulin distribution”, Fig. 1l). Our analysis and calculations showed that the ratio of stable to unstable MTs decreased for acetylated tubulin as a marker for stable MTs towards the neurite tip, while it increased for detyrosinated tubulin as a marker for stable MTs towards the neurite tip (Extended Data Fig. 1c).

To test which of the two ratios recapitulated the MT distributions, we measured the fraction of stable MT mass from the decay of MT mass after photoconversion in the neurite close to and far from the soma, at one third and two thirds of neurite length towards the neurite tip, respectively (Fig. 1m). The fraction of stable MT mass decreased from 34 ± 5% close to the soma to 23 ± 5% far from the soma (Fig. 1n-p), showing that the amount of stable MTs decreased towards the neurite tip relative to unstable MTs. This suggests that acetylated tubulin recapitulates stable MTs in developing neurites better than detyrosinated tubulin (Fig. 1j).

We further validated our results by directly comparing the photoconversion results to measured MT distributions by calculating the normalized change of the stable to unstable MT ratio from close to the soma to far from the soma (Fig. 1q, Extended Data Fig. 1d). This normalized change was similar to the value for acetylated tubulin but very different from detyrosinated tubulin as markers for stable MTs (-0.35 for photoconversion results, -0.25 for acetylated tubulin, 0.09 for detyrosinated tubulin) (Fig. 1q, Extended Data Fig. 1d). Therefore, we used non-acetylated and acetylated tubulin as markers for unstable and stable MTs, respectively, from here onwards (Fig. 1k,l). Taken together, the amount of unstable MTs remains relatively constant in the proximal half of the neurite and decreases distally, while the amount of stable MTs decreases along the whole neurite (Fig 1k,l). These data were the basis for uncovering how MT dynamic properties affect MT distributions in developing neurites.

### Developing a biophysical model of MT stabilization in neurites

We next investigated the role of MT dynamic properties in regulating MT distributions in developing neurites from a theoretical perspective. In mitotic cells, MT arrays during cell division are well defined through biophysical models that integrate several mechanisms such as basic MT dynamics known from *in vitro* data and modulation by motor proteins^27–33^. We hypothesized that a data-driven biophysical model will help to understand the development and maintenance of neuronal MT arrays. We therefore developed a one-dimensional model of unstable and stable MTs that undergo MT-RF (Fig. 2a-b; Methods section “Description of the core model”).

**Fig. 2.**
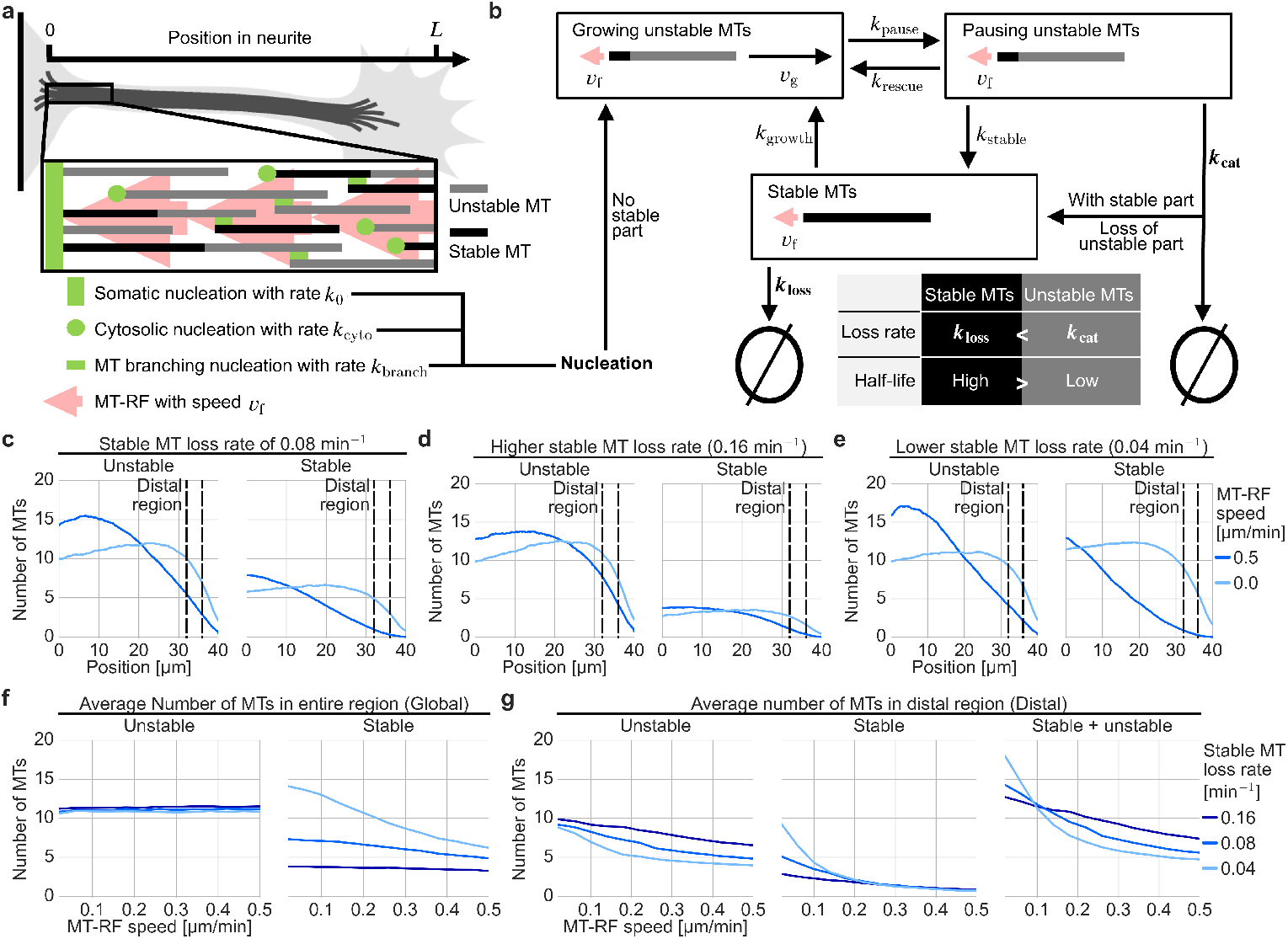
MT-RF reduces stable and distal MTs depending on the stable MT loss rate. **a**, Illustration of the three mechanisms for MT nucleation and MT-RF. **b**, Illustration of the states that MTs can be in, the transitions between these states and how new MTs can be generated and MTs can be lost through catastrophe. The illustration also shows the key difference of stable and unstable MTs in their loss rate and half-life. **c-g**, Simulations with different MT-RF speeds (0 – 0.5 µm/min) and different stable MT loss rates (0.04, 0.08, 0.16 min^-1^) (n = 1000 simulations per condition, simulation time: 450 min). (**c-e**) The distribution of unstable and stable MTs is plotted for simulations with fast MT-RF speed of 0.5 µm/min or without MT-RF (MT-RF speed of 0 µm/min) for different stable MT loss rates. Dashed lines indicate the region used to calculate distal MTs for the analysis in **g**. (**f-g**) The number of MTs averaged (**f**) in the entire region (global) or (**g**) from 80% (32 µm) to 90% (36 µm) region length towards the end of the region (distal) when changing the MT-RF speed for different stable MT loss rates.

Previous models of neuronal MTs lacked MT-RF nor included the difference between stable and unstable MTs^34–36^. To include stable and unstable MTs, our model characterizes these MTs with a long and short half-life, respectively, and describes the transitions between these states (Fig. 2b). To account for the local nucleation differences in each neuronal compartment^37,38^, we investigated local MT distributions by incorporating three known nucleation mechanisms in neurons. New MTs can be nucleated by somatic nucleation from MT organizing centers^37,39,40^, uniformly in the cytosol of the neurite^41,42^ or from the lattice of pre-existing MTs by MT branching^37,42^ (Fig. 2a); MT branching nucleation is limited by a global resource (Methods section “Description of the core model”). Importantly, MT-RF is modeled through all MTs moving retrogradely towards the soma at the same speed, consistent with our previous finding that the entire MT array moves uniformly in a retrograde fashion^16^ (Fig. 2a). Additionally, the model incorporates well known MT dynamics, including growth of unstable MTs from the tip of stable MTs and catastrophe^43,44^. The model contains 17 parameters, of which 7 parameters were measured here, calculated, or taken from literature (Table 1; Extended Data Fig. 2a-g). The remaining 10 parameters were estimated from the comparison to 6 experimental data points and an additional 3 experimental distributions (Table 1; Methods section “Fitting parameter values”). To obtain model MT distributions, we performed Monte Carlo simulations and numerically integrated the corresponding mean-field model (Extended Data Fig. 3a-c; Methods). These MT distributions provide the basis to investigate how known MT dynamics and MT-RF interact to influence stable and unstable MTs.

**Table 1.**
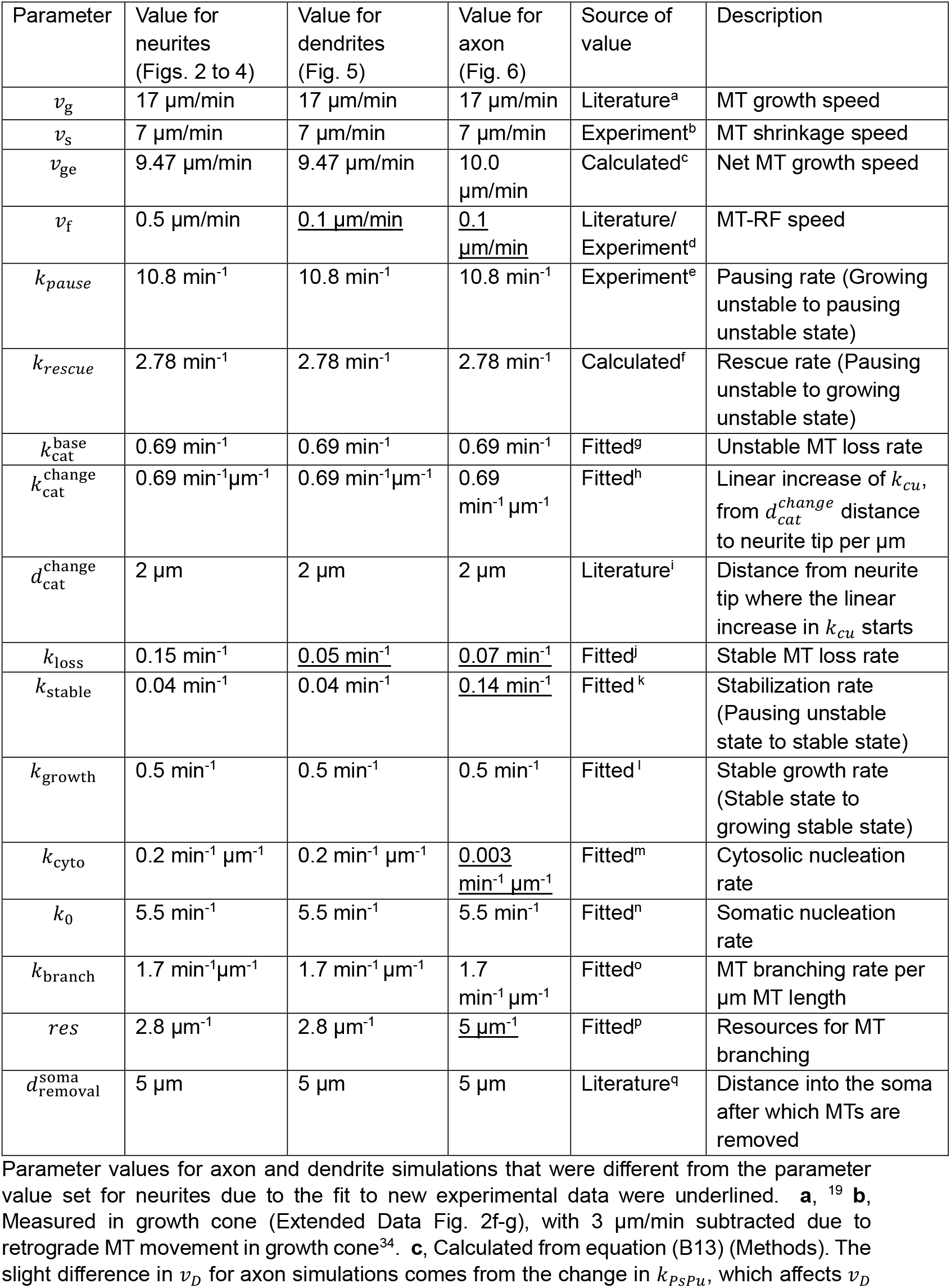
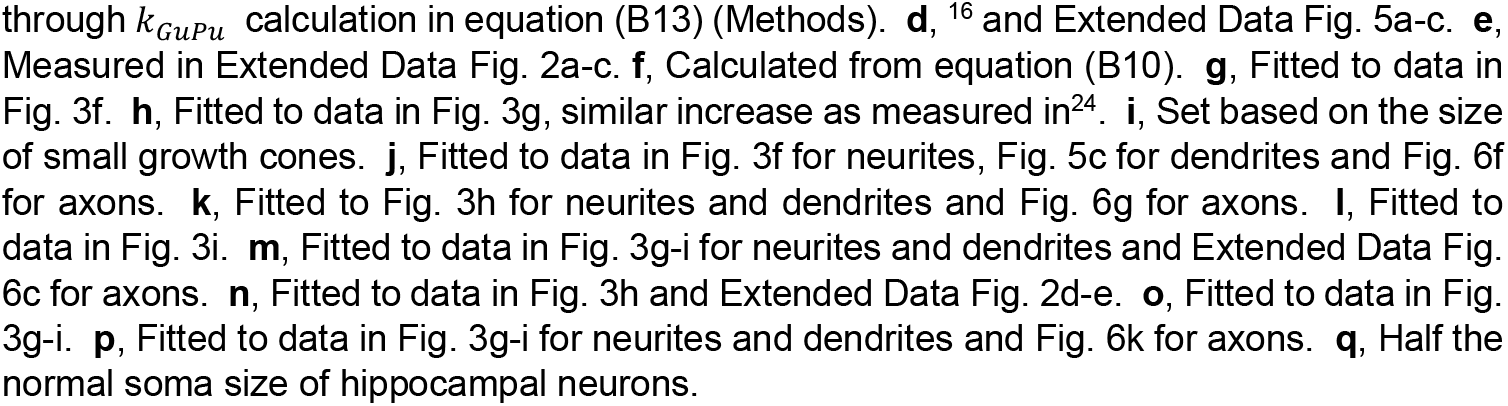
Standard parameter values used for key simulations.

| Parameter | Value for neurites (Figs. 2 to 4) | Value for dendrites (Fig. 5) | Value for axon (Fig. 6) | Source of value | Description |
| --- | --- | --- | --- | --- | --- |
| $v_g$ | 17 $\mu\text{m}/\text{min}$ | 17 $\mu\text{m}/\text{min}$ | 17 $\mu\text{m}/\text{min}$ | Literature<sup>a</sup> | MT growth speed |
| $v_s$ | 7 $\mu\text{m}/\text{min}$ | 7 $\mu\text{m}/\text{min}$ | 7 $\mu\text{m}/\text{min}$ | Experiment<sup>b</sup> | MT shrinkage speed |
| $v_{ge}$ | 9.47 $\mu\text{m}/\text{min}$ | 9.47 $\mu\text{m}/\text{min}$ | 10.0 $\mu\text{m}/\text{min}$ | Calculated<sup>c</sup> | Net MT growth speed |
| $v_f$ | 0.5 $\mu\text{m}/\text{min}$ | <u>0.1 $\mu\text{m}/\text{min}$</u> | <u>0.1 $\mu\text{m}/\text{min}$</u> | Literature/ Experiment<sup>d</sup> | MT-RF speed |
| $k_{\text{pause}}$ | 10.8 $\text{min}^{-1}$ | 10.8 $\text{min}^{-1}$ | 10.8 $\text{min}^{-1}$ | Experiment<sup>e</sup> | Pausing rate (Growing unstable to pausing unstable state) |
| $k_{\text{rescue}}$ | 2.78 $\text{min}^{-1}$ | 2.78 $\text{min}^{-1}$ | 2.78 $\text{min}^{-1}$ | Calculated<sup>f</sup> | Rescue rate (Pausing unstable to growing unstable state) |
| $k_{\text{cat}}^{\text{base}}$ | 0.69 $\text{min}^{-1}$ | 0.69 $\text{min}^{-1}$ | 0.69 $\text{min}^{-1}$ | Fitted<sup>g</sup> | Unstable MT loss rate |
| $k_{\text{cat}}^{\text{change}}$ | 0.69 $\text{min}^{-1}\mu\text{m}^{-1}$ | 0.69 $\text{min}^{-1}\mu\text{m}^{-1}$ | 0.69 $\text{min}^{-1}\mu\text{m}^{-1}$ | Fitted<sup>h</sup> | Linear increase of $k_{cu}$, from $d_{\text{cat}}^{\text{change}}$ distance to neurite tip per $\mu\text{m}$ |
| $d_{\text{cat}}^{\text{change}}$ | 2 $\mu\text{m}$ | 2 $\mu\text{m}$ | 2 $\mu\text{m}$ | Literature<sup>i</sup> | Distance from neurite tip where the linear increase in $k_{cu}$ starts |
| $k_{\text{loss}}$ | 0.15 $\text{min}^{-1}$ | <u>0.05 $\text{min}^{-1}$</u> | <u>0.07 $\text{min}^{-1}$</u> | Fitted<sup>j</sup> | Stable MT loss rate |
| $k_{\text{stable}}$ | 0.04 $\text{min}^{-1}$ | 0.04 $\text{min}^{-1}$ | <u>0.14 $\text{min}^{-1}$</u> | Fitted<sup>k</sup> | Stabilization rate (Pausing unstable state to stable state) |
| $k_{\text{growth}}$ | 0.5 $\text{min}^{-1}$ | 0.5 $\text{min}^{-1}$ | 0.5 $\text{min}^{-1}$ | Fitted<sup>l</sup> | Stable growth rate (Stable state to growing stable state) |
| $k_{\text{cyto}}$ | 0.2 $\text{min}^{-1}\mu\text{m}^{-1}$ | 0.2 $\text{min}^{-1}\mu\text{m}^{-1}$ | <u>0.003 $\text{min}^{-1}\mu\text{m}^{-1}$</u> | Fitted<sup>m</sup> | Cytosolic nucleation rate |
| $k_0$ | 5.5 $\text{min}^{-1}$ | 5.5 $\text{min}^{-1}$ | 5.5 $\text{min}^{-1}$ | Fitted<sup>n</sup> | Somatic nucleation rate |
| $k_{\text{branch}}$ | 1.7 $\text{min}^{-1}\mu\text{m}^{-1}$ | 1.7 $\text{min}^{-1}\mu\text{m}^{-1}$ | 1.7 $\text{min}^{-1}\mu\text{m}^{-1}$ | Fitted<sup>o</sup> | MT branching rate per $\mu\text{m}$ MT length |
| $res$ | 2.8 $\mu\text{m}^{-1}$ | 2.8 $\mu\text{m}^{-1}$ | <u>5 $\mu\text{m}^{-1}$</u> | Fitted<sup>p</sup> | Resources for MT branching |
| $d_{\text{removal}}^{\text{soma}}$ | 5 $\mu\text{m}$ | 5 $\mu\text{m}$ | 5 $\mu\text{m}$ | Literature<sup>q</sup> | Distance into the soma after which MTs are removed |

### MT-RF slowdown increases stable and distal MTs depending on the stable MT half-life

Since MT-RF slows down during neuronal development in the axon and later in dendrites^16,17^, we first explored the distributions of unstable and stable MTs with fast and slow MT-RF. The number of unstable MTs remains constant close to the soma (proximal) and then decreases towards the end (distal) for both the fast MT-RF of 0.5 µm/min measured in developing neurites^16^ and without MT-RF (MT-RF speed set to 0 µm/min) (Fig. 2c). In contrast, the number of stable MTs decreases across the entire region with fast MT-RF of 0.5 µm/min (Fig. 2c). Only without MT-RF, the number of stable MTs remains constant proximally and decreases distally, similar to unstable MTs (Fig. 2c). This suggests that MT-RF slowdown increases stable MTs more strongly than unstable MTs.

Since stable MTs are key for axon development^15^, we next asked why MT-RF slowdown is increasing stable MTs more strongly than unstable MTs. The key difference between stable and unstable MTs is that the reaction rate which removes stable MTs – the stable MT loss rate - is lower compared to the reaction rate at which unstable MTs are removed – the catastrophe rate (Fig. 2b). Thus, we hypothesized that increasing the stable MT loss rate results in a smaller increase of stable MTs upon MT-RF slowdown. Indeed, for a higher stable MT loss rate, MT-RF slowdown from 0.5 µm/min to 0 µm/min barely increases the average number of stable MTs in the entire region (global stable MTs, Fig. 2d,f). Conversely, for a lower stable MT loss rate, this MT-RF slowdown massively increases global stable MTs (Fig. 2e,f). Global unstable MTs are not changed by MT-RF for all stable MT loss rates (Fig. 2d-f). This shows that MT-RF slowdown increases stable MTs more than unstable MTs due to the lower stable MT loss rate. In other words, MT-RF moves longer lived MTs (which have a lower stable MT loss rate (Extended Data Fig. 4a,b)) further towards the soma during their longer lifetime, reducing them more. Consequently, MT-RF slowdown leads to a stronger increase for longer lived MTs.

For axon growth, in addition to stable MTs, MTs need to reach the distal tip^12,15,19,20^. We therefore investigated the distal MT contribution in relation to MT-RF and stable MT-RF loss rates. Interestingly, for all stable MT loss rates, MT-RF slowdown from 0.5 µm/min to 0 µm/min increases stable and unstable MTs distally more than proximally (Fig. 2c-e, Extended Data Fig. 4c). The corresponding increase of the average number of stable and unstable MTs in the distal region (distal MTs) by MT-RF slowdown is largest for a low stable MT loss rate (Fig. 2g), leading to a high overall increase in distal MTs (Fig. 2g, “Stable + unstable”). Thus, MT-RF slowdown not only increases stable MTs but also overall distal MTs; this increase becomes stronger for lower stable MT loss rates.

### MT dynamic properties explain experimental unstable and stable MT distributions

To test the predicted increase of stable and distal MTs by MT-RF slowdown, we first explored whether the interplay of modelled MT dynamics can explain experimental dynamics and distributions of stable and unstable MTs in neurons. Indeed, the model reproduces the experimental mass decay curve by adjusting only the catastrophe rate for unstable MTs and the loss rate of stable MTs while keeping the other parameters constant (Fig. 3a-d; Table 1; see Methods section “Statistics”). Using the same set of parameters (Table 1), the model reproduces the fraction of stable MT mass close to and far from the soma (Fig. 3e,f) and the distribution of stable and unstable MTs, except very close to the soma (Fig. 3g).

**Fig. 3.**
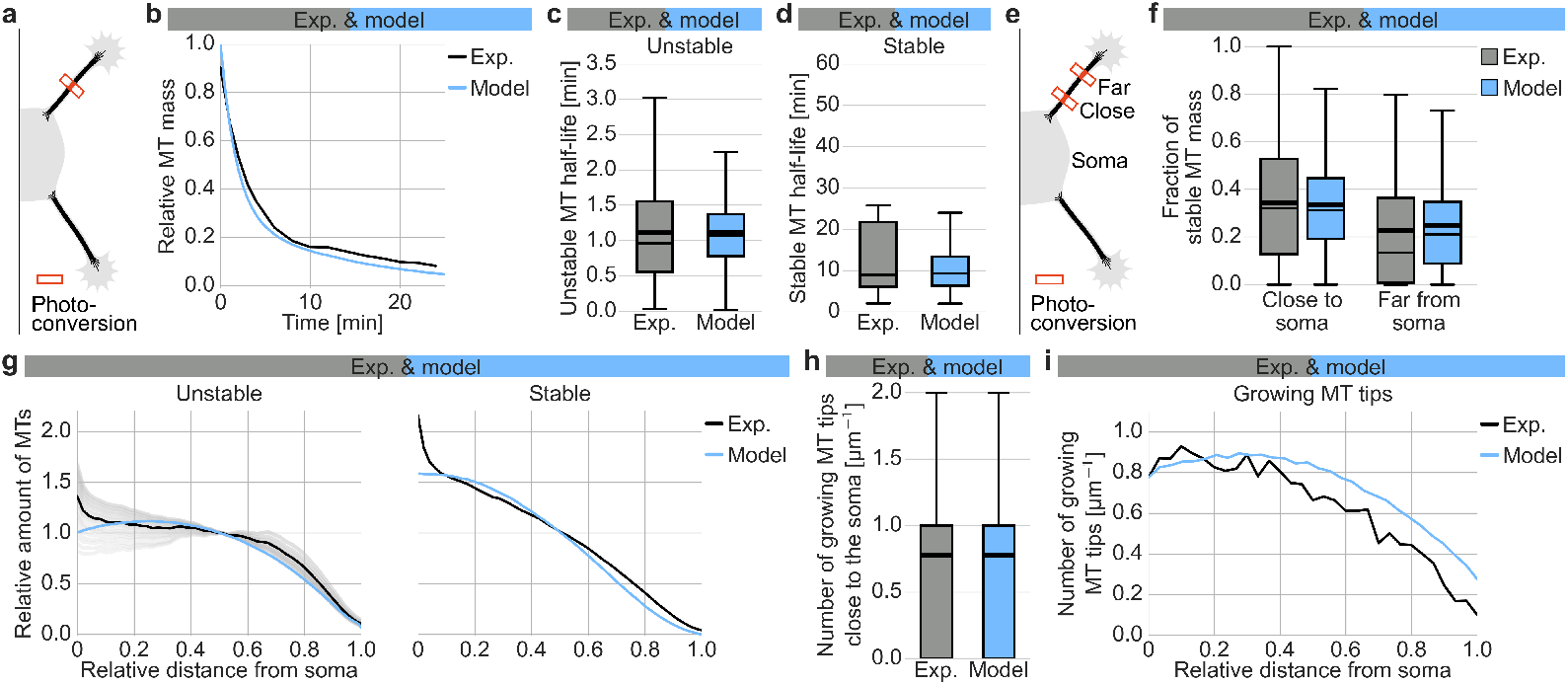
The model reproduces experimental data in developing neurites. **a-d**, Simulations of the photoconversion experiment over 40 µm are run and results compared to experimental data from Fig. 1, d-f. The total photoconverted MT mass was measured at all timepoints and the decay curve fitted to a dual exponential function. (**a**) Illustration of photoconversion experiment (model for b-d: 1000 simulations, simulation time: 510 min). (**b**) Average MT mass decay curve (n = 43 neurons, N = 4 experiments). (**c-d**) The half-life of (**c**) the fast-decaying component (unstable MTs) or (**d**) the slow-decaying component (stable MTs) (Exp.: **c**: n = 43 neurons, N = 4 experiments; **d**: n = 40 neurons, N = 4 experiments). **e-f**, Simulations of photoconversion close to and far from the soma over 40 µm are run and results are compared to experimental data from Fig. 1p. (**e**) Illustration of photoconversion experiment. (**f**) The fraction of stable MT mass close to and far from the soma was calculated from the dual exponential fit of the slow and fast decaying component (n = 1000 simulations, simulation time: 510 min). **g**, Distribution of stable and unstable MTs for simulations over 30 µm and experiments from Fig. 1k-l with an average neurite length of 30 µm. For simulation data, the distribution of stable MTs and unstable MTs is summed for all MT populations. Values are normalized to the value in the middle of the neurite (Exp: Stable (acetylated): n = 505 neurites from 143 neurons, N = 3 experiments; Model: n = 16000 simulations, simulation time: 450 min). **h,i**, The density of growing MT tips was calculated from the data of EB3 experiments from ^16^ with an average neurite length of 30 µm. The density of growing MT tips is calculated from simulations over 30 µm (n = 37 neurons, N = 3 experiments; **g-i**: n = 16000 simulations, simulation time: 450 min). The density (**h**) close to the soma or (**i**) across the entire neurite was plotted. Due to the large number of simulations, model and experimental data are not statistically compared (Methods section “Statistics”). The thick line in boxplots indicates the mean, the thin line the median.

For additional experimental data to compare the model to, we obtained the experimental distribution of growing MT tips from live cell imaging data of fluorescently labeled end-binding protein 3 (EB3)^16^. Indeed, the experimental density of growing MT tips close to the soma matched the density of growing tips in the model (Fig. 3h). Further, the experimental distribution of growing MT tips remained relatively constant in the first 40% of the neurite and then decreased towards the neurite tip (Fig. 3i), which is fitted well by the model (Fig. 3i). In summary, our comparative analyses of experimental data and model predictions suggest that the modelled MT dynamics explain distributions of unstable and stable MTs as well as growing MT tips in developing neurites. This indicates that the model could be useful for making predictions about how MT-RF affects stable and unstable MTs.

### MT-RF slowdown increases stable MTs distally in developing neurites

Since MT-RF slows down early in axon development^16^, we investigated how slow MT-RF in young axons affects stable and unstable MTs. Keeping the same set of parameters (Table 1), the model predicts that changing the MT-RF speed from 0.5 µm/min to 0.2 µm/min, the speed in young axons after one day in culture^16^, increases stable MTs by 21.8% globally but not unstable MTs (Fig. 4a,b). Interestingly, in the distal region, the model predicts that both stable and unstable MTs increase noticeably with slow MT-RF, by 148.7% and 35.0%, respectively (Fig. 4b).

**Fig. 4.**
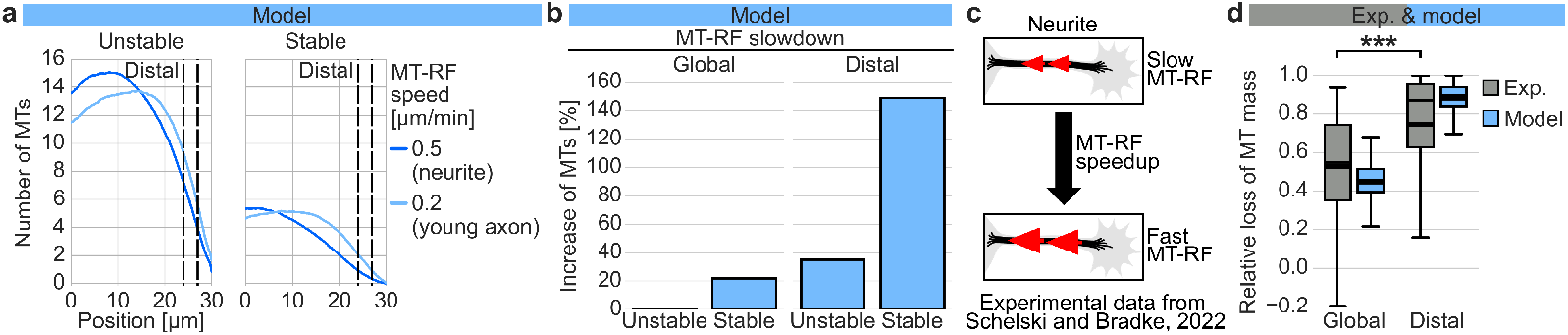
MT-RF slowdown in young axons increases stable MTs in the distal neurite. **a,b**, Simulations over 40 µm are run with the fast MT-RF speed of developing neurites (0.5 µm/min) or the slow MT-RF speed of young, 1-day old axons (0.2 µm/min; n = 2000 simulations, simulation time: 450 min). (**a**) The distributions of stable and unstable MTs summed from all populations are quantified with the region used to calculate distal MTs in **b** marked by dashed lines. (**b**) The average number of stable and unstable MTs in the entire neurite (global) or from 32 µm (80%) to 36 µm (90%; distal) is calculated. The increase in MTs for slow MT-RF of 0.2 µm/min compared to fast MT-RF of 0.5 µm/min is calculated. **c,d**, Analyzed experimental raw data and simulations of the MT-RF speedup experiment from ^16^ (raw data underlying Fig. 10, A to F). (**c**) Illustration of MT-RF speedup experiment in **d**. (**d**) The loss of MTs was calculated as the difference of the MT mass before MT-RF speedup to the lowest MT mass up to 15 min after the start of MT-RF speedup. MT mass loss was calculated in the entire neurite (global) or from 60% to 90% neurite length towards the neurite tip. Simulations are run over 32 µm, the average length of a neurite in the experimental data. For simulations, MT-RF speed was increased from 0.2 µm/min to 2.41 µm/min after 450 min (when the steady state was reached), with these values being the experimentally measured average MT-RF speeds before and after speedup (Experiment: n = 59 neurites from 21 cells, N = 4 experiments; Simulation: n = 500, simulation time: 549 min; p = 3.8*10^-7^, Dunn’s test. Due to the large number of simulations, model and experimental data are not statistically compared and thus only statistically significant differences between experimental data are shown (Methods section “Statistics”). The thick line in boxplots indicates the mean, the thin line the median.

To validate the predicted stronger effect of MT-RF on distal MTs in developing neurites, we compared model predictions with our published experimental data for the speedup of MT-RF in developing neurites using membrane-recruited dynein^16^ (Fig. 4c). Using this experimental data for MT-RF speedup to 2.41 µm/min ± 0.15 µm/min^16^ we measured the corresponding loss of MTs. For this MT-RF speedup and using the stable MT loss rate matching the experimental stable MT half-life of approx. 10 min (Fig. 1e), the model predicts an MT loss of 44.7% ± 0.4% globally and a higher MT loss of 88.0% ± 0.4% in the distal region (Fig. 4d). Thus, our theoretical prediction can provide an explanation for the experimental data, where the MT loss was 52.5% ± 3.8% in the entire neurite and also higher in the distal neurite with 74.4% ± 4.0% (Fig. 4d). This shows that the model can account for the experimentally measured MT loss by only changing MT-RF to experimentally measured values while keeping other parameters unchanged. It suggests that the model recapitulates the experimental effect of MT-RF on MTs in developing neurites. In summary, MT-RF slowdown in developing neurites mainly increases stable MTs, particularly in the distal neurite.

### MT-RF slowdown in dendrites increases stable MTs globally and distally

In dendrites, stable MTs are crucial for cargo sorting^9^. Yet, it is unclear whether stable MTs increase in dendrites during development. Additionally, MT-RF slows down in dendrites to 0.1 µm/min, albeit later than in axons^16^. Since we found that MT-RF slowdown increases stable MTs in developing neurites (Fig. 4), we tested whether stable MTs also increase in dendrites, and whether this is fueled by MT-RF slowdown. Interestingly, in addition to slow MT-RF, we found experimentally that the half-life of stable MTs is longer in dendrites than in developing neurites (Fig. 5a,b). We lower the stable MT loss rate in the model to match this higher experimental stable MT half-life (Fig. 5c). To model MT dynamics in the dendrite, we use this lower stable MT loss rate and slower MT-RF speed but otherwise keep the same set of parameters as for developing neurites (Table 1). The model predicts that these two changes together allow dendrites to increase the amount of stable MTs globally by 203.6% and distally by 408.8%, compared to the scenario with higher stable MT loss rate and faster MT-RF, as in neurites (Fig. 5d,e). This stable MT increase results in a stable MT mass fraction of 59.3% ± 0.7% in the middle of the dendrite, which is similar to the experimentally measured value of 57.8% ± 4.8% (Fig. 5f). This shows that dendrites massively increase stable MTs, transport roads for dendritic cargo ^9^, by only slowing down MT-RF and lowering the stable MT loss rate.

**Fig. 5.**
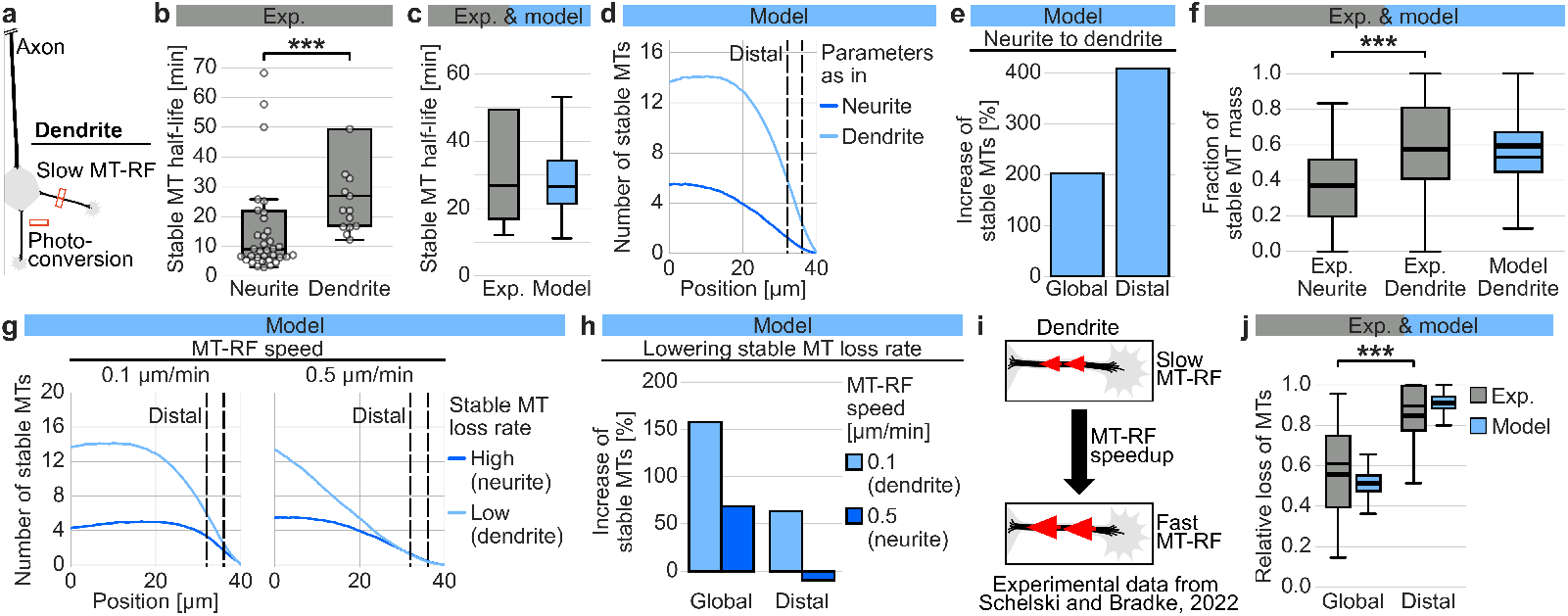
MT-RF slowdown in dendrites increases stable MTs globally and distally. **a,b**, Neurons cultured for six to seven days and expressing the tubulin TUBB2a fused to the photoconvertible fluorophore mEos3.2 were imaged after a small MT patch was photoconverted in dendrites, at an average 46% neurite length towards the neurite tip (Raw data from Fig. S7^16^). The photoconverted MT mass was measured at each timepoint and the decay fitted to a dual exponential function. (**a**) Illustration of the photoconversion experiment in dendrites and slow MT-RF as the main parameter change used to simulate dendrites. (**b**) The half-life of stable MTs was calculated from the fitted rate of the slow decaying component. For neurites and dendrites, three and four datapoints with a half-life above 500 min are out of the plotting range, respectively. (Dendrite: n = 17 neurons, N = 1 experiment; Data from neurites from Fig. 1e: n = 38 neurons, N = 4 experiments; p = 0.0004, Mann-Whitney U test) **c**, Simulations with the measured slow dendritic MT-RF speed of 0.1 µm/min were run and the stable MT loss rate was reduced (from 0.15 min^-1^ to 0.05 min^-1^) to obtain the experimental stable MT half-life from **b** (n = 1000 simulations, simulation time: 540 min). **d,e**, Simulations over 40 µm are run with the fast fitted stable MT loss rate (Fig. 3d) and fast MT-RF speed of 0.5 µm/min from developing neurites (Neurite) or the slow fitted stable MT loss rate (fitted in **c**) and slow MT-RF speed of 0.1 µm/min from dendrites (Dendrite) (n = 2000 simulations, simulation time: 450 min). (**d**) The distribution of stable MTs is plotted with the region used to calculate distal MTs in (e) marked by dashed lines. (**e**) The average number of MTs in the entire region (global) or from 32 µm (80%) to 36 µm (90%; distal) is calculated. The increase in stable MTs for parameter values simulating the dendrite compared to parameter values simulating the neurite is calculated (n = 500 simulations, simulation time: 350 min). **f**, The fraction of stable MT mass from experimental data of neurites at 30% neurite length towards the neurite tip (Fig. 1f; Exp neurite) and of dendrites at 46% dendrite length towards the neurite tip (Exp. Dendrite) were compared to simulations with parameter values corresponding to experimental data from dendrites (Exp. neurite data from Fig. 1f: n = 45 neurons, N = 4 experiments; Exp dendrite: n = 18 neurons, N = 1 experiment; p = 9.4*10^-5^, Dunn’s test; Simulation: n = 1000 simulations, simulation time: 540 min). **g,h**, Simulations over 40 µm are run with combinations of dendritic slow MT-RF (0.1 µm/min) and fast MT-RF of developing neurites (0.5 µm/min) with the high (0.15 min^-1^) and low (0.05 min^-1^) stable MT loss rate in developing neurites and dendrites, respectively (n = 2000 simulations, simulation time: 450 min). (**g**) Stable MT distributions are shown with the region used to calculate distal MTs in **h** marked by dashed lines. (**h**) MTs in the entire region (global) or from 32 µm (80%) to 36 µm (90%; distal) are analyzed. Then the increase in stable MTs for slower stable MT loss rate (0.05 min^-1^) compared to fast stable MT loss rate (0.15 min^-1^) is calculated. **i,j**, Simulations and analyzed raw experimental data of the MT-RF speedup experiment in dendrites from ^16^ (raw data underlying Fig. S12). (**i**) Illustration of speedup experiment. (**j**) Raw experimental data was from ^16^ and simulation data is analyzed right before dimerization and on average 50 min to 60 min after MT-RF speedup, either in the entire neurite (global) and from 60% to 90% neurite length towards the neurite tip (distal). Simulations were run over 42 µm, the average dendrite length in experimental data. For simulations, MT-RF speed was increased from 0.05 µm/min to 1.32 µm/min after 450 min (when the steady state was reached), these values being the experimentally measured average MT-RF speeds before and after speedup (raw data underlying ^16^, Fig. S12; Exp: n = 17 neurons, N = 3 experiments; Sim: n = 500 simulations, simulation time: 570 min; p = 0.00097, Dunn’s test). The thick line in boxplots indicates the mean, the thin line the median. Due to the large number of simulations, model and experimental data are not statistically compared and thus only statistically significant differences between experimental data are shown (Methods section “Statistics”).

To dissect the mechanisms underlying the increase in stable MTs, we tested the influence of MT-RF slowdown and low stable MT loss rate separately. The model predicts that with slow MT-RF of 0.1 µm/min, lowering the stable MT loss rate increases the amount of stable MTs globally and distally, by 158.0% and 63.5%, respectively (Fig 5g,h). In contrast, with fast MT-RF of 0.5 µm/min, lowering the stable MT loss rate increases the amount of stable MTs only by 69.0% globally and reduces distal stable MTs by 9.5% (Fig. 5g,h). Thus, these results suggest that dendrites can maintain a high amount of stable MTs globally and distally by synergistically reducing the stable MT loss rate and slowing down MT-RF.

We validated the predicted effect of MT-RF on MTs in dendrites by comparing model predictions with our published experimental data for MT-RF speedup in dendrites using membrane-recruited dynein^16^ (Fig. 5i). Using these experimental data with MT-RF speedup to 1.39 µm/min ± 0.3 um/min^16^ we measured the corresponding MT loss in dendrites. For this MT-RF speedup and using the same parameter values as in dendrites (Table 1), the model predicts a global MT loss after MT-RF speedup of 51.2 % ± 0.2% and a higher distal MT loss of 91.5% ± 0.2% (Fig. 5j). This matches the experimental global MT loss of 55.7% ± 7.0% and the higher experimental distal MT loss of 84.6% ± 4.3% (Fig. 5j). Hence, it demonstrates that changing only MT-RF speed to experimentally measured values can explain the experimentally measured MT loss. Further, it shows that the model makes validated predictions about dendrites by only changing the two parameters stable MT loss rate and MT-RF speed while keeping the other parameters used to model developing neurites unchanged (Table 1). In summary, with slow MT-RF, increasing stable MT half-life leads to more global and distal stable MTs. In contrast, with faster MT-RF this increase of global stable MTs is reduced and the increase of distal stable MTs reverses to a decrease. Thus, dendrites need to slow down MT-RF in addition to increasing stable MT half-life to maintain a high amount of stable MTs.

### MT-RF slowdown in axons allows MTs to increase towards the axon tip

Slowdown of MT-RF is a key step for axon development^16,17^. Additionally, stable MTs are known to be crucial for axons, with MT stabilization being sufficient for axon development^15^. In particular, distal MTs are crucial for fast axonal growth^12,15,19,20^. The consistent increase of stable and distal MTs by MT-RF slowdown in developing neurites and dendrites prompted us to investigate whether MT-RF slowdown also increases stable and distal MTs in axons. We first tested whether the model can help to understand the formation of the MT array in axons. Axons differ from developing neurites and dendrites^3,20,45^. We focus on the following two key differences (Fig. 6a): longer shaft length of 160 µm, compared to 30-40 µm used for developing neurites and dendrites, and lower cytosolic nucleation rate^42^. We also measured three additional differences experimentally: 0.1 µm/min slow MT-RF (Extended Data Fig. 5a-c), longer stable MT half-life (Fig. 6b-d) and a higher fraction of stable MT mass (Fig. 6e). These five differences are implemented in the model by changing five parameters, while keeping the other parameters as in developing neurites and without adding additional mechanisms to the model (Table 1). To fit the model to the longer stable MT half-life and higher fraction of stable MT mass, we lower the stable MT loss rate and increase the stabilization rate (Fig. 6f,g). The lower cytosolic nucleation rate is key to obtain a model that is in line with the uniform plus-end-out MT orientation in axons^5^ (Extended Data Fig. 6a-c). Specifically, MT branching allows newly nucleated MTs to have the same orientation as the MT from which they were nucleated^46,47^ (Extended Data Fig. 6a) and thereby does not directly determine MT orientation. Conversely, nucleation from the soma leads to plus-end-out MTs (Extended Data Fig. 6a). Lastly, MTs nucleated in the cytosol of the axon have an equal chance to be plus-end-out or plus-end-in (Extended Data Fig. 6a), which reduces the percentage of plus-end-out MTs in axons^42,46^. This suggests that cytosolic MT nucleation is kept low in the axon to allow uniform plus-end-out MT orientation^42^. Without a reduction of the cytosolic nucleation rate, the percentage of plus-end-in MTs in the model is 43.3 ± 0.1% and thus much higher than the experimentally measured value of 2.5% (Extended Data Fig. 6b). To obtain a realistic level of cytosolic nucleation rate for the axon, we reduce this nucleation rate to achieve a percentage of plus-end-in MTs of 2.8%, close to experimentally measured value of 2.5%^5^ (Extended Data Fig 6c).

**Fig. 6.**
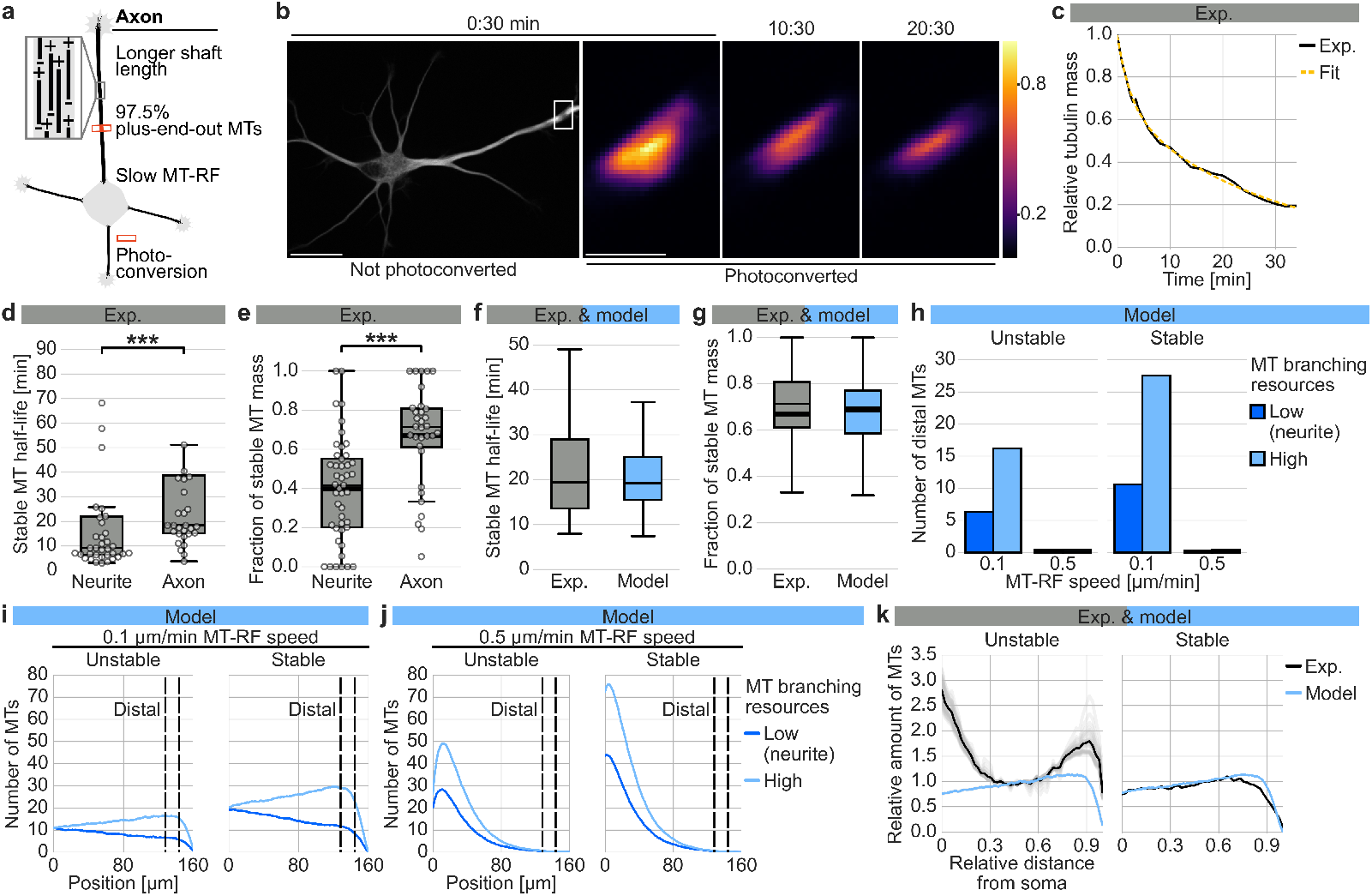
MT-RF slowdown in axons enables stable MTs to increase towards the axon tip. **a-e**, Neurons expressing the tubulin TUBB2a fused to the photoconvertible fluorophore mEos3.2 were cultured for three days. Then a small MT patch in the axon was photoconverted at an average 32% neurite length towards the axon tip and the summed photoconverted intensity measured for each timepoint. (**a**) Illustration of photoconversion experiments and differences of axons compared to neurites that were implemented in simulations. (**b**) Representative neuron for **d,e**. Intensity is color-coded from purple (low) to yellow (high). Time is in min:seconds. Scale bars, 20 µm for not zoomed and 4 µm for zoomed images. (**c**) Decay curve of summed photoconverted intensity with fitted dual exponential function of representative neuron in **b**. (**d-e**) Decay curves were fitted to dual exponential function and (**d**) half-life and (**e**) fraction of stable MT mass were plotted for neurites and axons. For neurites and axons in **d**, three and six datapoints with a half-life above 10,000 min are out of the plotting range, respectively. (neurite data from Fig. 1,e-f; **d**: neurite: n = 38 neurons, axon: n = 32 neurons, p=0.00099; **e**: neurite: n = 45 neurons, axon: n = 32 neurons, p = 1.7 * 10^-5^; N = 4 experiments for all groups, Mann-Whitney U test). **f,g**, Comparison of experimental data from **d** and **e** and simulations over 160 µm for (**f**) stable MT half-life and (**g**) fraction of stable MT mass obtained from fitted dual exponential curve (n = 500 simulations, simulation time: 1660 min). To fit simulation results to experimental data, the stable MT loss rate is lowered (from 0.15 min^-1^ to 0.07 min^-1^) and stabilization rate is increased (from 0.04 min^-1^ to 0.14 min^-1^). **h-j**, Simulations with (**i**) slow MT-RF in 3-day-old axons of 0.1 µm/min and (**j**) fast MT-RF of developing neurites of 0.5 µm/min and with high (5.0 µm^-1^) or low (2.8 µm^-1^, as in simulations of developing neurites and dendrites) resources for MT branching (n = 500 simulations, simulation time: 1600 min). (**h**) Average number of distal MTs from simulations in **i,j** from 128 µm (80%) to 144 µm (90%). (**i,j**) Distribution of unstable and stable MTs with the region used to calculate distal MTs marked by dashed lines. **k**, Comparison of the distribution of stable and unstable MTs from experiments and simulations from **i** with a high amount of MT branching resources. Values were normalized to the middle of the axon. For the experimental distributions, neurons cultured for three days were stained for tyrosinated and acetylated or tyrosinated and detyrosinated tubulin and the corresponding distributions were measured for 100 µm to 200 µm long axons. Intensities were quantified for each point along the axon by summing up all pixel intensities across the axon width at that point and normalizing the value by the point closest to the soma. The distribution of non-acetylated tubulin was calculated as in Fig. 1l and used as the distribution for unstable MTs, while the distribution of acetylated tubulin was used as the distribution for stable MTs (Acetylated: n = 56 neurons; N = 3 experiments). The thick line in boxplots indicates the mean, the thin line the median. Due to the large number of simulations, model and experimental data are not statistically compared and thus only statistically significant differences between experimental data are shown (Methods section “Statistics”).

Since MT-RF slows down in axons compared to developing neurites, we first tested the effect of slow MT-RF compared to fast MT-RF on stable and unstable MT distributions. With the parameter changes noted above and slow MT-RF of 0.1 µm/min, the model predicts that the number of stable and unstable MTs decreases towards the distal region (Fig. 6i; low resources for MT branching). Since MT branching is more prevalent in axons than in developing neurites^42,46^ and since a reduction of MT branching impairs axon development^42^, we tested the effect of increased MT branching. In our model, MT branching is mainly limited by the amount of available resources. With higher amounts of resources for MT branching, but keeping other parameters unchanged, the number of stable and unstable MTs increases towards the axon tip for slow MT-RF of 0.1 µm/min (Fig. 6h,i; high resources for MT branching). In contrast, with fast MT-RF of 0.5 µm/min, the number of stable and unstable MTs decreases towards the axon tip even with higher amounts of resources for MT branching (Fig. 6h,j). This suggests that MT-RF slowdown allows the higher MT branching in axons^42,46^ to increase distal MTs (Fig. 6h). Thus, MT-RF slowdown in axons not only directly increases distal MTs but also enables the increase of distal MTs by increased MT branching.

We next wanted to test our prediction that stable MTs in axons increase towards the axon tip. To this end, we stained neurons with an older axon after three days in culture for tyrosinated tubulin as a marker for younger MTs as well as acetylated and detyrosinated tubulin as markers for older MTs (Extended Data Fig. 7a-e). As for developing neurites (Fig. 1g-l, Extended Data Fig. 1a-j), we used the distribution of acetylated tubulin as a marker for stable MTs and calculated the distribution of non-acetylated tubulin to obtain the distribution of unstable MTs (Extended Data Fig. 7a-e). The resulting distributions for stable and unstable MTs matched the change of the ratio of stable to unstable MTs from close to the soma to far from the soma, measured by live cell imaging of MT decay (Extended Data Fig. 7f-i). The amount of unstable MTs decreases in the proximal half of the axon and then increases (Fig. 6k). Importantly, the amount of stable MTs increases until close to the axon tip (Fig. 6k). The unstable MT distribution cannot be fit well by the model (Fig. 6k), particularly close to the soma, arguing for additional molecular mechanisms that play a role in this region, such as the emerging axonal initial segment (AIS)^45,48^. Notwithstanding, the distribution of stable MTs is fit well by the model throughout the axon (Fig. 6k). This supports the prediction that distal stable MTs are increased in axons through a combination of MT-RF slowdown and increased MT branching (Fig. 6h-j).

### MT-RF slowdown maintains distal MTs in axons while allowing high stable MT half-life and uniform plus-end-out MTs

Since distal MTs are required for fast axonal growth^12,15,19,20^, we investigated the mechanisms regulating distal MTs in axons. First, we characterized the role of axonal MT-RF slowdown for distal MTs by performing simulations with different MT-RF speeds (Fig. 7a). The model predicts that without MT-RF slowdown in the axon, using the MT-RF speed of 0.5 µm/min in developing neurites, barely any distal unstable or stable MTs are left (Fig. 7b-d), leaving the distal axon without MT: the MT array is only 0.7 ± 0.0 MTs thick (Fig. 7d). Only with MT-RF speeds below 0.3 µm/min, some distal MTs remain (Fig. 7d). Further slowdown from the speed in young axons of 0.2 µm/min to the speed in old axons of 0.1 µm/min is also relevant since it increases distal MTs two-fold (Fig. 7d). This suggests that MT-RF slowdown is needed to maintain distal MTs in axons. While the increase of distal MTs by MT-RF slowdown in axons (Fig. 7b,c) is consistent with our results in developing neurites (Fig. 4) and dendrites (Fig. 5), it is much stronger in axons. Therefore, axons are particularly dependent on MT-RF slowdown to maintain distal MTs. This raises the question of which axon-specific mechanisms lead to the low number of distal MTs without MT-RF slowdown.

**Fig. 7.**
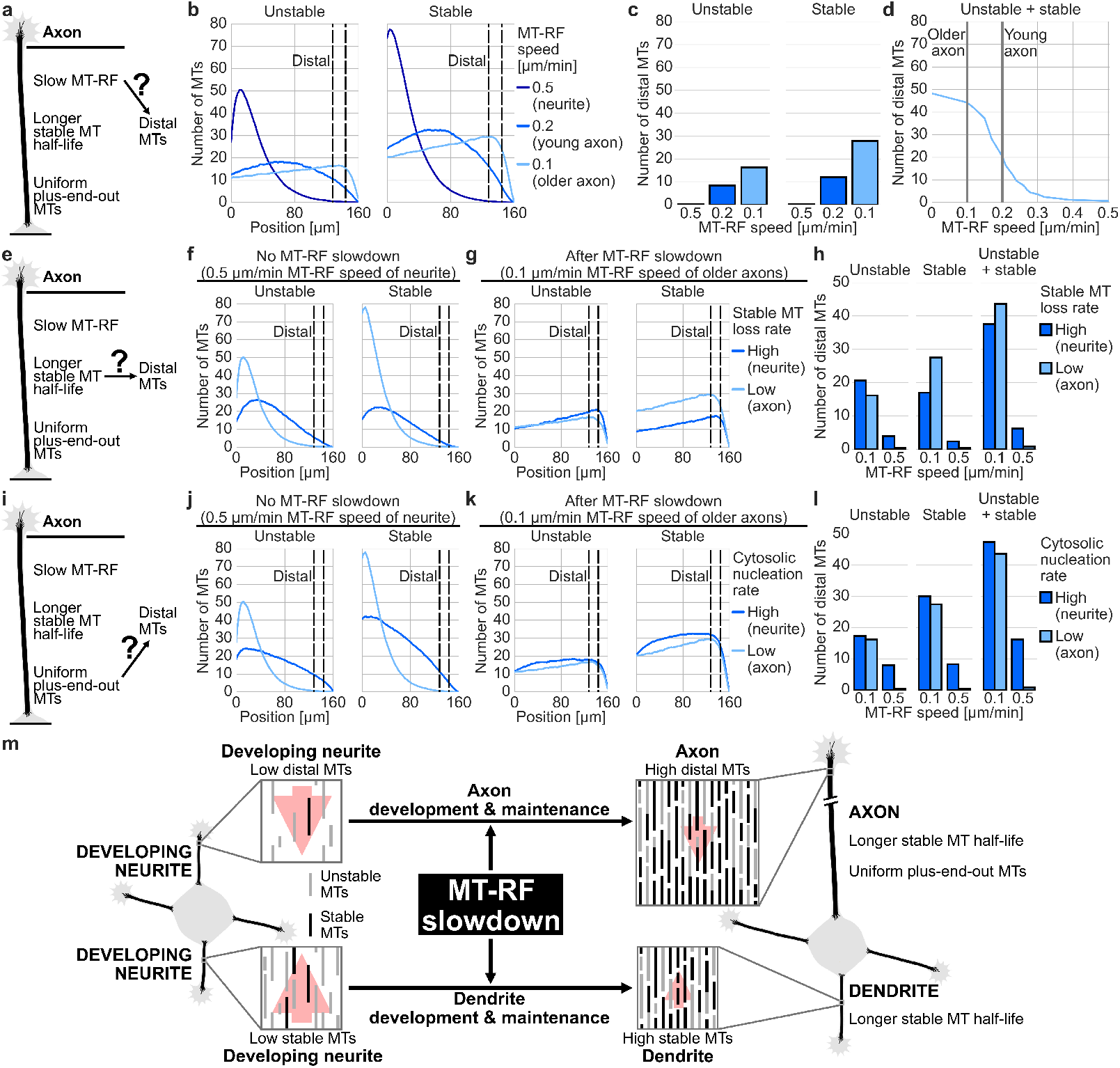
MT-RF slowdown is needed for distal MTs due to axonal MT properties. **a-l**, Stable, unstable and the sum of stable and unstable MTs are quantified from simulations over 160 µm (**b,f,g,j,k**) across the entire region. Dashed lines in **b,f,g,j,k** indicate the distal region from 128 µm (80%) to 144 µm (90%) used to calculate distal MTs for the analyses in **c,d,h,l**. (**a-d**) Simulations with MT-RF speed from developing neurites (0.5 µm/min), young axons (0.2 µm/min) or 3-day-old axons (older axons, 0.1 µm/min) (n = 500 simulations, simulation time: 1600 min). (**a**) Changes implemented in the axon based on experimental data. Slow MT-RF could influence distal MTs in the axon. (**b**) Distribution of unstable and stable MTs. (**c**) The average number of distal unstable and stable MTs for different MT-RF speeds. (**d**) The average number of distal MTs depending on the MT-RF speed. MT-RF speed of 1-day-old axons (young axon; 0.2 µm/min) and 3-day-old axons (older axon, 0.1 µm/min) are marked by grey lines. (**e-h**) Simulations with combinations of the fast stable MT loss rate of developing neurites (0.15 min^-1^) or the slow stable MT loss rate of axons (0.07 min^-1^) with slow axonal MT-RF (0.1 µm/min) or fast MT-RF (0.5 µm/min) (n = 500 simulations, simulation time: 1600 min). (**e**) Slow stable MT loss rate could influence distal MTs in the axon. (**f,g**) Distribution of unstable and stable MTs for (**f**) fast MT-RF and (**g**) slow MT-RF. (**h**) The average number of stable, unstable and total distal MTs for fast and slow stable MT loss rate. (**i-l**) Simulations with the low cytosolic nucleation rate fitted to the fraction of plus-end-in MTs in Extended Data Fig. 6a-c (0.003 min^-1^µm^-1^) or the high cytosolic nucleation rate as used in developing neurites and dendrites (Figs. 2-5; 0.2 min^-1^µm^-1^), combined with slow MT-RF of 0.1 µm/min or fast MT-RF of 0.5 µm/min were run (n = 500 simulations, simulation time: 1600 min). (**i**) The high proportion of plus-end out MTs could influence distal MTs in the axon. (**j,k**) Distribution of stable and unstable MTs for (**j**) fast MT-RF and (**k**) slow MT-RF. (**l**) The average number of stable, unstable and total distal MTs. **m**, MT-RF slowdown increases distal MTs in the axon due to the increased effect and stable MTs in dendrites to drive the development of both axons and dendrites.

Since stable MTs are key for axonal development^15^, reducing the stable MT loss rate to increase stable MTs is an important mechanism for axons. Therefore, we second tested the role of the lower stable MT loss rate, and thus the longer stable MT half-life (Fig. 6f) in axons (Fig. 7e), compared to the higher stable MT loss rate in developing neurites (Figs. 3-4). The model predicts that without MT-RF slowdown, lowering the stable MT loss rate to the axonal value massively reduces both distal unstable and distal stable MTs, so that barely any distal MTs are left (Fig. 7f-h). In contrast, with the slow MT-RF speed of 0.1 µm/min in old axons, lowering the stable MT loss rate hardly reduces distal unstable MTs and even increases distal stable MTs (Fig. 7g,h). Thus, lowering the stable MT loss rate in axons requires MT-RF slowdown to prevent loss of distal MTs. This also shows that a lower stable MT loss rate is not only a key mechanism to increase stable MTs but also allows MT-RF to tightly regulate distal axonal MTs.

Third, we tested the role of uniform plus-end-out MT orientation (Fig. 7i), which is a key property of axons to enable axonal transport and cargo sorting^8^. To investigate the role of uniform plus-end-out MT orientation, we tested the effect of lowering the cytosolic nucleation rate from the value used in developing neurites and dendrites (Figs. 3-5) to the much lower value fitted to the high plus-end-out MT fraction in axons (Fig. 6). Lowering the cytosolic nucleation rate without MT-RF slowdown massively reduces distal MTs, from 16.2 ± 0.2 MTs to 0.8 ± 0.1 MTs (Fig. 7j-l). For slow MT-RF, lowering the cytosolic nucleation rate does not change distal MTs noticeably (Fig. 7j-l). This suggests that uniform plus-end-out MT orientation makes MT-RF slowdown necessary to retain distal MTs and thereby maintain axonal length. In conclusion, the model suggests that slowdown of MT-RF together with its interplay with two axon-specific changes is important for maintaining long axons and fast axon growth through retaining distal MTs. In other words, MT-RF needs to slow down in axons to maintain distal MTs, while also keeping a high fraction of stable MTs and uniform plus-end-out MTs.

In summary, by integrating experimental data on MT dynamics and MT distributions in a biophysical model we uncover the central role of MT-RF slowdown for axons and dendrites (Fig. 7m). In dendrites, MT-RF slowdown enables a large increase in stable MTs. In axons, MT-RF slowdown is necessary for MTs to reach the axon tip. The strong effects of MT-RF slowdown in dendrites and axons are caused by increased stable MT half-life; in the axon, this effect is even further augmented due to the uniform MT orientation. The interaction of MT-RF slowdown with these two MT properties and MT branching is sufficient to explain the changes in the stable MT distribution for axon and dendrite development.

## Discussion

So far, axon and dendrite development have been thought to require different mechanisms in axons and dendrites to trigger morphological and molecular polarization^2^. Our data overturn this view by showing that the same mechanism, slowdown of MT-RF, enables development and maintenance of the axon and dendrites. By investigating the distribution of stable and unstable MTs we reveal how distinct MT dynamics in axons and dendrites drive changes in stable MTs, required for polarized transport, and distal MTs that promote axon growth. We find that the interaction of MT-RF with only three MT dynamics - MT stability, cytosolic MT nucleation and MT branching - is sufficient to explain the change of the stable MT distribution for both axon and dendrite development. Our unifying perspective leads to four unexpected insights.

First, neurite growth is thought to increase through the regulation of distal MTs by the growth cone^49,50^. Here, we find that in addition distal MTs are strongly regulated by MT-RF acting on the entire MT array to redistribute MTs away from the neurite tip. This redistribution reduces neurite growth, which is supported by MT-RF speedup inducing neurite retraction^16^. After MT-RF slowdown, increased MT branching in axons^42^ allows axons to increase their distal MTs. Interestingly, reduced MT branching through HAUS7 knockout reduces axon regeneration^51^. This emphasizes a new avenue of looking beyond growth cone-mediated mechanisms towards axon shaft-mediated mechanisms that can redistribute MTs towards the growth cone to increase axon growth.

Second, while axons are known to require an increase in stable MTs^15^, dendrites are mainly characterized as having fewer stable MTs than axons^52–54^. While the fraction of stable MTs increases among minus-end-out oriented MTs during dendrite development^9,55^, it has remained unexplored whether overall stable MTs increase or whether stable MTs only change their orientation^55^. We now find that dendrites massively increase stable MTs. Previously, knockout of the MT stabilizing protein MAP2 in Purkinje cells led to a 23% loss of MT density in dendrites^14^. While this was discounted as being a minor decrease^56^, it matches our predicted MT loss of 28% for decreasing the dendritic stable MT half-life to the level of developing neurites and in fact corresponds to a large predicted stable MT loss of 60%. This supports the role of increased stable MTs in dendrites and highlights the insights gained when comparing experimental results with model predictions.

Third, it has remained unclear which cell-inherent mechanism allows dendrites to switch from the non-growing state of developing neurites to a growing state. Interestingly, KO of the MT stabilizing protein MAP2 reduces dendritic length^14^, suggesting that stable MTs are needed for dendrite growth, as known in axons. Nonetheless, it was thought that a potential trigger for dendritic growth would be based on different mechanisms than axonal growth^57^. Instead, we found that the same mechanism that triggers stable axon growth could also allow dendrites to grow through increasing distal MTs and stable MTs: slowdown of MT-RF. This further highlights the unifying features of our model centering around the MT-RF.

Lastly, intuitively one would assume MT stabilization should be sufficient to increase stable MTs for axon and dendrite development. However, we find that MT-RF suppresses the stabilization-mediated increase in stable MTs by reducing stable MTs with a longer half-life even more. Thus, MT-RF slowdown is needed in addition to MT stabilization for developmentally relevant increases in stable MTs. This can also explain how MT-RF slowdown in the whole neuron directs Kinesin-1 to all neurites and subsequently leads to the development of multiple axons^17^. Before axon development, stable MTs stochastically increase in single neurites, probably mediated by increasing stable MT half-life, which recruits the MT motor protein Kinesin-1 into the neurite through more detyrosinated MTs^24,58,59^. However, with fast MT-RF, the increase in stable MTs - driven by their extended half-life - is attenuated, which could explain its transient nature in developing neurites^58^. In contrast, MT-RF slowdown allows a steady increase in stable MTs, stabilizing Kinesin-1 recruitment and axon identity.

Why does early MT-RF slowdown lead to axon development, whereas later it leads to dendrite development? The key difference is the orientation of MTs that are stabilized. Axonal cargo is steered into axons through detyrosinated, stable MTs which are plus-end-out oriented^59^. Thus, if plus-end-out MTs are stabilized, then an axon forms. In dendrites on the other hand stable MTs keep axonal cargo out, through changing their orientation to plus-end-in^9,55^. Therefore, if stabilized MTs change their orientation to plus-end-in then a dendrite forms.

Intriguingly, tuning of MT flow could be a conserved mechanism to regulate length depending on MT half-life across different biological contexts. During cell division, MTs move towards the poles of the mitotic spindle at around 0.6 µm/min^60^, which slows down in anaphase^61,62^ to enable spindle elongation^63^. This is remarkably similar to MT-RF in neurons, which moves MTs retrogradely at 0.5 µm/min and slows down in axons and dendrites^16,17^, which we found in this study to enable axon and dendrite elongation. As in neuronal development, also during cell division, the half-life of MTs changes in different phases, which regulates chromosomal instability in cancer cells^64^. MT half-life in anaphase can even become as high as in dendrites^62^. Thus, the half-life dependent effect of MT-RF we found in neurons could also be relevant for MT flux during cell division. Thus, our model with MT stabilization could be employed to investigate the effect of MT stability during cell division.

Importantly, the MT cytoskeleton is perturbed in several neurodevelopmental diseases^65–67^, neurodegenerative diseases^68–70^ and spinal cord injury^71,72^, often showing MT loss, changes in MT stability or changes in tubulin acetylation. However, which changes in MT dynamics are sufficient to cause these pathological phenotypes has remained unclear. Interestingly, for several neurodevelopmental^66^ and neurodegenerative diseases^73^, as well as spinal cord injury^18,74^, mild MT stabilization improves the condition. While the similarities we uncovered between axons and dendrites suggest that MTs could be pathologically changed in both axons and dendrites, our data also highlight that the same treatment may lead to different outcomes in axons and dendrites. Our data-driven biophysical model would allow to systematically characterize which experimentally measured changes in MT dynamics are sufficient to explain pathological changes in MT distributions in axons and dendrites. This could provide mechanistic avenues towards finding combinations of MT dynamics that can be changed to combat disease.

### Online Methods

#### Description of the core model

We developed a one-dimensional model of MTs in the neurite, where the origin of the coordinate system is located at the soma and the neurite tip is at *x* = *L*. Each MT consists of a stable and unstable part of lengths *l*_*s*_ and *l*_*u*_, respectively. The position of the MT, *x*, represents the position of the minus-end of the MT, while the sum of the position *x*, the stable length *l*_*s*_, and the unstable length *l*_*u*_ represents the position of the plus-tip of the MT. Based on the dynamic properties of MTs, the model describes three key populations of MTs: those that are (i) unstable and growing, (ii) unstable and pausing, and (iii) stable (Fig. 2a-b). In the model, all MTs have their plus-end oriented towards the neurite tip (plus-end-out), while in neurons only 80% of MTs are oriented in this way^5^. MT stabilization is modeled as a stabilization of the MT tip and thus is a transition from an unstable MT to a stable MT.

To describe how the density of unstable growing MTs plus ends changes in time, *t*, we describe their dynamics by a transport equation:

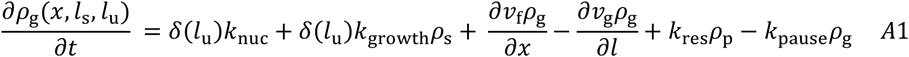

Here, flow velocity, growth velocity, pausing rate (transition from the growing to the pausing state) and rescue rate (transition from the pausing to the growing state) are denoted *v*_f_, *v*_g_, *k*_pause_ and *k*_res_, respectively. The growth rate is the transition from the stable state to the growing state and thus represents the growth of an unstable MT from the tip of a stable MT and is denoted *k*_growth_. This leads to the formation of well-known MTs that are stable towards the minus-end and unstable towards the plus-end^43,44^ (Fig. 2b). The nucleation rate, 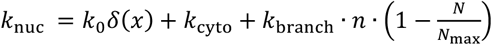 includes nucleation from the soma with rate *k*_0_, from the cytosol with rate *k*_cyto_, and along pre-existing MTs with rate *k*_branch_. The number of microtubules along the neurite is given as 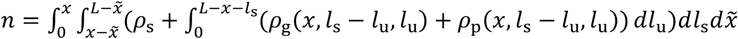, and the total number of microtubules in the neurite is 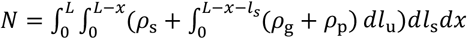. The number of available resources for nucleation along pre-existing MTs is denoted *N*_max_. *δ*is the dirac delta function.

To describe how the density of unstable pausing MT plus ends changes in time t, we describe their density by another transport equation:

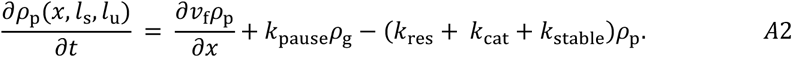

The catastrophe rate (loss of the unstable length of the MT *l*_*u*_) and stabilization rate (the transition from the pausing unstable state to the stable state) are denoted *k*_cat_ and *k*_stable_, respectively. The other parameters are described above.

Lastly, to describe how the density of stable MT plus ends changes in time t, we describe their density by the following equation:

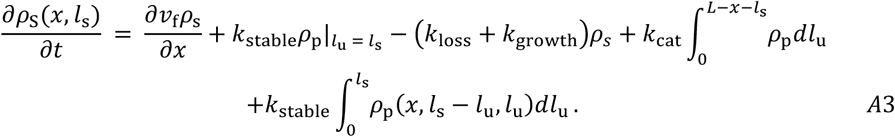

The loss rate (complete loss of the stable MT) is denoted as *k*_loss_. The other parameters are as described above. The integral for the catastrophe of pausing unstable MTs 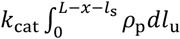 represents the loss of the unstable length *l*_u_ of the pausing unstable MT *ρ*_p_, after which only the stable length *l*_s_ remains. Thus, the density *ρ*_p_ is integrated across all MTs with the same stable length *l*_s_ at position *x*. The integral for the stabilization of pausing unstable MTs 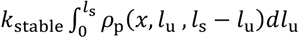 represents the stabilization of the unstable length *l*_u_ of the pausing unstable MT *ρ*_p_ to stable length *l*_s_. Thus, the density *ρ*_p_ is integrated along the line of MTs with the same total length *l*_s_ + *l*_u_ with their minus end at the same position *x*.

This set of equations represent the core of the model. Additionally, changes at the boundary of the neurite tip have to be taken into account.

#### Analytical mean-field solutions

We first solve a simplified system of partial differential equations (Extended Data Fig. 3b), with flow velocity *v*_f_, nucleation from the soma *k*_0_ and nucleation from preexisting MTs *k*_branch_ set to zero. We also set the growth rate *k*_growth_ to zero. As a consequence, there is no growth of unstable MTs from the tip of stable MTs and unstable MTs do not have a stable length.

The change in the density of growing unstable MT plus-ends over time is described by the following transport equation:

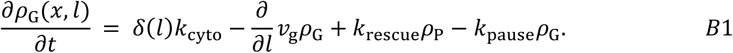

The nucleation rate from the cytosol, MT growth velocity, the rescue rate, and pausing rate are denoted as *v*_g_, *k*_rescue_, *k*_pause_, and *k*_cyto_, respectively. Here, *δ*is the dirac delta function.

The change in the density of pausing unstable MT plus-ends over time is described by the following equation:

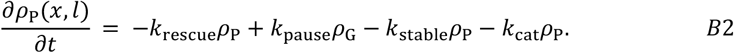

The stabilization and catastrophe rate are denoted as *k*_stable_ and *k*_cat_, respectively.

The change in the density of stable MT plus-ends over time is described by the following equation:

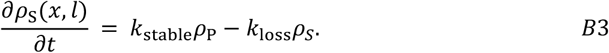

The loss rate is denoted as *k*_loss_.

To solve the steady state, all derivatives with respect to time are set to zero:

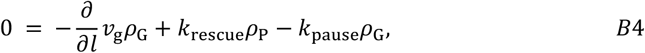

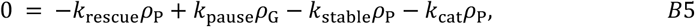

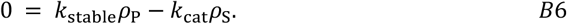

Solving the algebraic equation (B5) leads to the solution for *ρ*_P_:

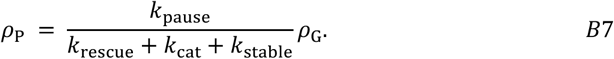

Solving the algebraic equation (B6) leads to the solution for *ρ*_*S*_:

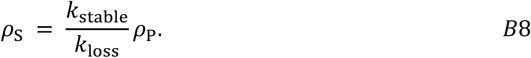

Plugging equation (B7) in equation (B4) leads to:

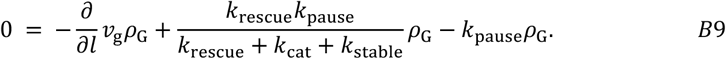

The general solution of this first order ordinary differential equation is:

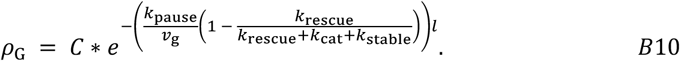

To obtain the boundary condition at *l* = 0, the steady state ODE is integrated around zero:

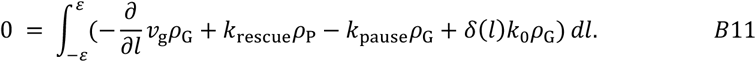

This shows that:

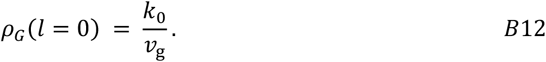

Consequently, the steady state solution for *ρ*_G_ is:

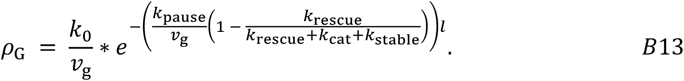

#### Numerical mean-field solutions

To numerically solve the steady state of the two systems of partial differential equations in Extended Data Fig. 3a,c, a custom implementation of the finite difference algorithm with a forward Euler method is used. The code is implemented using the Python package numba, to just-in-time compile the code to run fast on the central processing unit (CPU).

For Extended Data Fig. 3a we set both the nucleation from the soma *k*_0_ and nucleation from preexisting MTs *k*_*branch*_ to zero. Then we solve the steady state of the following system of partial differential equations, also accounting for differences at the boundary of the neurite tip.

The change in the density of unstable MT plus ends changes in time t is described by the following transport equation:

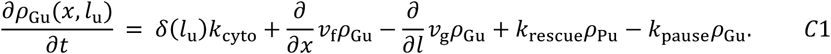

The nucleation rate from the cytosol, flow velocity, MT growth velocity, rescue rate and pausing rate are denoted as *k*_cyto_, *v*_f_, *v*_g_, *k*_rescue_ and *k*_pause_, respectively. Once again, *δ*represents the dirac delta function.

The change in the density of pausing unstable MT plus-ends over time t is described by the following transport equation:

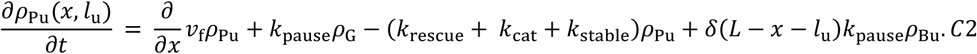

The catastrophe rate and stabilization rate and denoted as *k*_*cat*_ and *k*_*stable*_, respectively.

At the boundary of the neurite tip, the change of the density of unstable growing MT plus-tips over time t is described by the following transport equation:

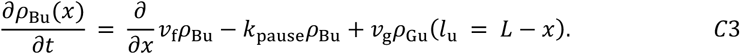

The change in the density of plus-ends of stable MTs over time *t* is described by the following transport equation:

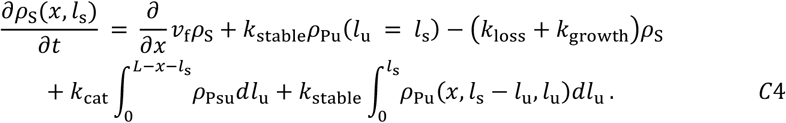

The change in the density of unstable MT plus ends growing from stable MTs over time t is described by the following transport equation:

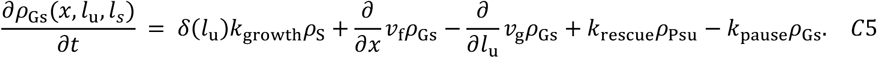

The change in the density of growing unstable MT plus ends growing from stable MTs over time t is described by the following transport equation:

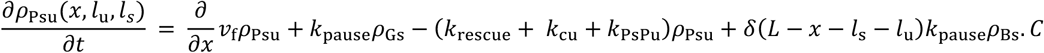

At the boundary of the neurite tip, the change of the density of unstable growing MT plus-tips growing from stable MTs over time t is described by the following transport equation:

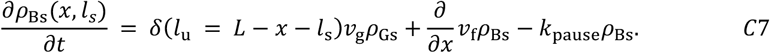

We then solved the steady state of a simplified system of partial differential equations (Extended Data Fig. 3c) numerically with the nucleation rate from pre-existing MTs *k*_branch_ not set to zero but no resource limit for this nucleation. However, we set flow velocity *v*_*f*_ and nucleation from the soma *k*_0_ to zero. We also set the growth rate *k*_growth_ to zero. As a consequence, there is no growth of unstable MTs from the tip of stable MTs and unstable MTs do not have a stable length.

The change in the density of growing unstable MT plus ends changes in time t, is described by the following transport equation:

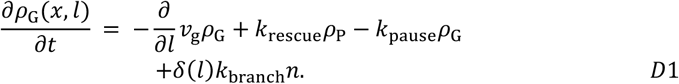

The MT growth velocity, rescue rate, pausing rate and nucleation rate from preexisting MTs are denoted as *v*_g_, *k*_rescue_, *k*_pause_, and *k*_branch_. The density of pausing unstable MT plus-ends and stable MT plus-ends are denoted as *ρ*_P_ and *ρ*_S_. *δ*is the dirac delta function. The number of microtubules along the neurite is given as 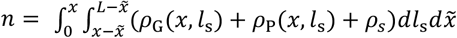.

The change in the density of pausing unstable MT plus ends changes in time t, is described by the following equation:

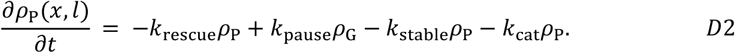

The stabilization rate and catastrophe rate are denoted as *k*_stable_ and *k*_cat_.

The change in the density of stable MT plus ends changes in time t, is described by the following equation:

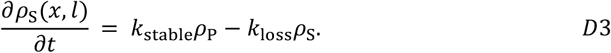

The loss rate is denoted as *k*_loss_.

### Running stochastic simulations

MTs in neurites are simulated with a defined dimension, corresponding to the neurite length. Position *x* = 0 corresponds to the start of the neurite at the soma and *x* = *L* corresponds to the neurite tip. MTs are not allowed to extend beyond the neurite tip, but the minus-end is allowed to move out of the neurite. MTs are simulated with a property for the position of the minus end, one property for its unstable and one property for its stable length. MTs are removed from the simulation through MT-RF reducing their position property, when their plus-end would be more than 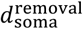 out of the neurite, which was set at 5 µm and thus a position lower than -5. This distance represents MTs being completely depolymerized when reaching the middle of the soma. Once MTs are removed from that position, if they were nucleated by MT branching, the corresponding resource is available again.

Stochastic simulations are run with a custom-implementation of the exact next reaction Stochastic Simulation Algorithm (SSA)^75^ . The algorithm is implemented using the Python package numba, to just-in-time compile the code to run many simulations in parallel on a graphics processing unit (GPU). A Python package with an application programming interface (API) was written to make stochastic simulations on GPUs easily useable (https://github.com/maxschelski/cytostoch). Importantly, due to approximate calculations of MT density for MT branching and of the position-dependent unstable MT loss rate (see below), simulations of the complete system are not exact. Parameter values are fitted by manually comparing simulation results to experimental data.

Due to the importance of MTs close to the neurite tip^12,15,19,20^ the neurite tip is explicitly modeled by increasing the unstable MT loss rate close to the neurite tip^24^. The increased position-dependent unstable MT loss rate is calculated as follows:

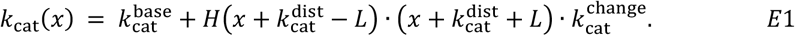

The baseline catastrophe rate, the change of the catastrophe rate per µm distance from the tip, and the maximum distance from the tip for which the catastrophe rate should be changed are denoted as 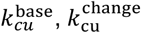 and 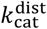. This results in an unstable MT loss rate that is constant throughout the neurite and linearly increases in the last micrometer to its maximum. The Heaviside function is denoted as *H*.

The total MT branching rate is calculated based on the following equation:

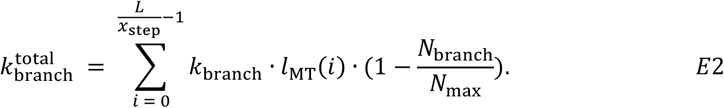

With *i* being the number of the step in *x*, and *x*_step_ is the step size, set as 1 µm (for 30 µm and 40 µm long neurites) up to 4 µm (for 160 µm long neurites). The rate for nucleation from pre-existing MTs is denoted as *k*_branch_. The total MT length from the *i* step to the *i* + 1 step is denoted as *l*_MT_(*i*) and is calculated by summing the length of all MTs in this *x* range. The total number of MTs nucleated by branching and the maximum number of MTs nucleated by branching are denoted by *N*_branch_ and *N*_max_, respectively.

Since MT length and MT position are changing, the reaction rate 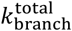 is a function of time *t* and thus the rate *λ* is a function of time *t* as well:

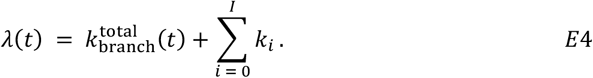

The sum of all reaction rates other than branching nucleation rate is denoted as 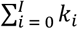, with *I* being the total number of other reaction rates.

Thus, the following integral needs to be solved to obtain Tau:

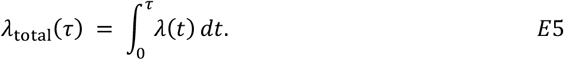

Tau is obtained as a random number, *R* drawn from an exponential distribution, divided by the total rate *λ*_*total*_:

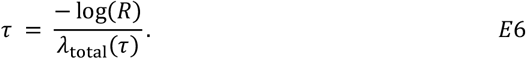

Due to the nonlinear Heaviside step functions due to absolute limits on MT growth, this integral is solved by iteratively obtaining better guesses of Tau. The first guess of Tau is based on the current sum of all rates, ignoring the time dependence. For each guess, the integral equation is solved, obtaining a value for the rate *λ*_*total*_ (*τ*) and thus a new value for Tau. The guess is therefore adjusted by moving it a bit closer to the new Tau. This is done for a maximum of 10 iterations. After obtaining the time until the next reaction tau, the next reaction executed after time tau is chosen according to the standard next-reaction Gillespie algorithm.

To identify the simulation time after which steady state is reached, simulation results with a time interval of 50 min are examined by eye. The timepoint after which no noticeable difference in the distributions is observed by eye was chosen as the simulation time. This simulation is generally the same between different simulations of neurites / dendrites / axons of the same length (e.g. for all simulations of 40 µm the same simulation time is used). Simulations of 30 µm neurites are generally run for 400 min, of 40 µm neurites and dendrites for 450 min, of 160 µm axons for 1600 min. The exact simulation times are noted for each panel in the figure legends.

Results for simulations of the MT-RF speedup experiment (in developing neurites and dendrites) are run until the steady state is reached, after which MT-RF speed is increased and data are extracted with a time interval of 1.5 min, which is the same interval as for experiments.

Results for simulations of MT decay experiments were extracted with a time interval of 0.5 min for up to 60 min (developing neurites and dendrites) or 90 min (dendrites and axons). The time interval matched experimental imaging times.

#### Mapping parameter values of growing-shrinking model to growing-pausing model

Switching of MTs between growing and shrinking^27,34^ is represented as a simplified switch between growing and pausing and the complete loss of the MT (catastrophe) (Fig. 2B). To still fit our model to the distribution of growing MT tips and the total number of MTs at the base of the neurite, the ratio of the number of growing MT tips and the total number of MTs in our growing-pausing model needed to be the same as in the standard biophysical growing-shrinking model for MTs. In the growing-shrinking model this ratio is determined by the shrinkage rate (the transition from the growing to the shrinking state) and the rescue rate (the transition from the shrinking to the growing state). To obtain a realistic ratio of growing MTs to total MTs, we use the shrinkage rate of the growing-shrinking model as pausing rate in our growing-pausing model and keep the same rescue rate as for the growing-shrinking model.

MTs in the growing-shrinking model shrink in the shrinking state but do not change their length in our growing-pausing model. Thus, the growth velocity in our growing-pausing model needs to account for the shrinking in the growing-shrinking model by calculating an effective growth velocity *v*_g_, which is lower than the actual MT growth velocity *v*_g_. This effective growth velocity is calculated by matching the expected length of the growing-shrinking and growing-pausing model.

The growing-shrinking model is governed by the following two partial differential equations for the density of growing MT tips *ρ*_*G*_ and shrinking MT tips *ρ*_*S*_:

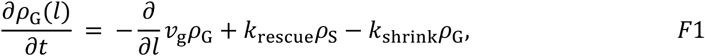

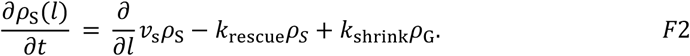

With *v*_g_ being the MT growth speed and *v*_*s*_ the MT shrinkage speed. The general steady state solution of *ρ*_G_ is:

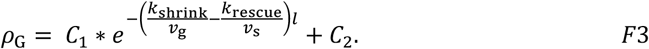

Thus, the expected length is:

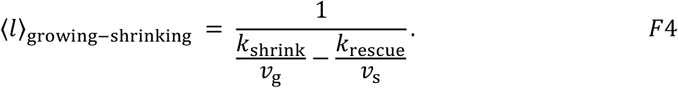

The lifetime of MTs *τ* is approximately the time it takes for the average MT length ⟨*l*⟩ to depolymerize plus the time it takes to depolymerize the MT length *l*_*τ*_ growing during the lifetime *τ*:

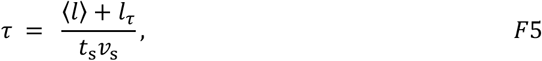

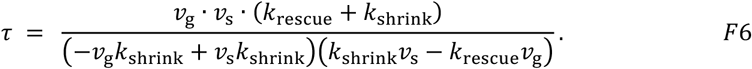

The MT length *l*_*τ*_ growing during the lifetime *τ* is calculated as:

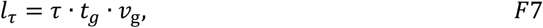

with *t*_*S*_ being the fraction of time a MT shrinks:

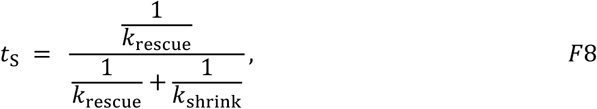

and with *t*_*G*_ being the fraction of time a MT grows:

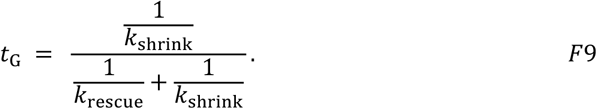

To calculate the rescue rate *k*_*GS*_, the following quadratic equation must be solved:

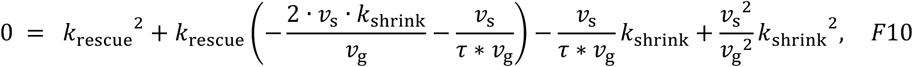

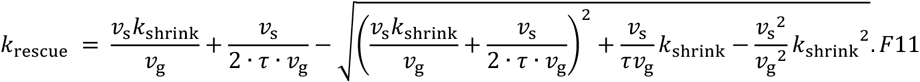

To solve this equation, we measured the shrinkage rate *k*_shrink_ from the average lifetime of growing MT tips in living neurons, visualized through fluorescently labeled end-binding protein 3 (EB3; Extended Data Fig. 2a-c)^16^. The calculated rescue rate *k*_rescue_ is used as the rescue rate for the growing pausing model. The measured shrinkage rate *k*_shrink_ is used as the pausing rate *k*_pause_ for the growing-pausing model:

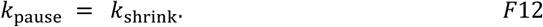

With the pausing rate *k*_pause_ and the rescue rate *k*_rescue_ set, the effective growth velocity *v*_ge_ of the growing-pausing model is calculated so that the expected length matches that of the growing-shrinking model. This effective growth velocity is lower than the actual MT growth speed to account for MTs shrinking.

The expected length of the growing-pausing model is:

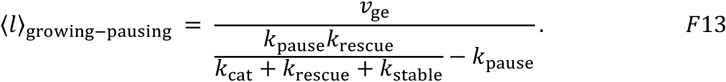

Setting eq. F13 equal to F4 and solving for the effective growth velocity, *v*_d_ yields:

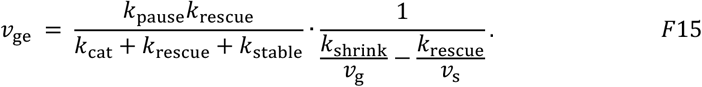

#### Fitting parameter values

To fit parameters, model results were compared to experimental data. Single or multiple parameters were varied to fit specific experimental data as described in Table 1. Parameter values were modulated manually, until the model results matched experimental data reasonably well.

#### Animals

All animal experiments were done in accordance with the Animal Welfare Act as well as the guidelines of the North Rhine-Westphalia State Environment Agency [Landesamt für Natur, Umwelt und Verbraucherschutz (LANUV); reference number for approval (Aktenzeichen: 84-02.04.2015.A334 and Aktenzeichen 81-02.04.2021.A208)]. Animals were housed in groups (up to five mice per cage) with controlled room temperature at 21° to 22°C in addition to an artificial 12-hour light:12-hour dark cycle (lights off at 6:00 p.m.). Mice (adult C57Bl6/J) were given food and water ad libitum throughout the experiment.

#### Neuron culture

Primary hippocampal neurons were dissected from embryonic day 16.5 (E16.5) to E17.5 mouse brains. Following dissection, hippocampi were collected in Hanks’ balanced salt solution with magnesium and calcium supplemented with 7 mM HEPES (Thermo Fisher Scientific, 14-025-050 and 15-630-056). Next, hippocampi were digested in 0.25% trypsin-EDTA supplemented with 7 mM HEPES (Thermo Fisher Scientific, 25-200-056) at 37°C for 15 min. Subsequently, they washed three times with minimum essential medium (MEM)–1% horse serum (MEM–1% HS) medium (1× MEM, 1× essential and nonessential amino acids, 2 mM l-glutamine (all from Thermo Fisher Scientific, 11-430-030, 11-130-036, 11-140-035, 25-030-081), 0.22% NaHCO3,

0.6% glucose, and 1% HS (Biowest, S0910-500)). Finally, hippocampi were mechanically dissociated with fire-polished glass-Pasteur pipettes. Coverslips were coated with poly-L-Lysine (1 mg/ml) (Sigma-Aldrich, P2636) in borate buffer at room temperature overnight, three to four times washed with double-distilled H_2_O (ddH_2_O) and placed in the incubator at 36.5°C and 5% CO_2_ with MEM-HS 10% to equilibrate for at least 2 hours before plating. Neurons were cultured at 36.5°C and 5% CO2 with a glia-feeding layer for staining neurons and without a glia-feeding layer for live-cell-imaging, as previously described^76^.

For fixating and staining neurons, 14 neurons/mm^2^ were plated on 15 mm glass-coverslips with self-made paraffin dots. Two to four hours after plating, glass coverslips with neurons were flipped on glia feeding layer with neuronal medium (N2) (1x MEM, 1 mM sodium pyruvate (Sigma-Aldrich, S8636), 1% Neuropan 2 supplement (Pan-Biotech, P07-11100), NaHCO_3_, 0.6% glucose, 2 mM L-glutamine and 1x B-27^™^ supplements (Thermo Fisher Scientific, 17-504-044) which was preconditioned for 3 days. For the glia feeding layer, 9 mouse glia cells/mm^2^ were plated in a 6 cm culture dishes and kept in MEM-HS10%. Three days before culture of neurons, media was exchanged to N2 for preconditioning.

For live-cell imaging of neurons, after transfection, 2 * 10^4^ neurons were plated in eight-well glass-bottom dishes (ibidi, 80827-90). Two to three hours after plating, MEM–10% HS medium was replaced by neuronal medium (N2) (1×MEM, 1 mM sodium pyruvate, 1% Neuropan 2 supplement (Pan-Biotech, P07-11100), NaHCO3, 0.6% glucose, 2 mM L-glutamine and 1xB-27^™^ supplements (Thermo Fisher Scientific, 17-504-044)), which was preconditioned by incubation on glia cells for two to three days.

#### Neuron transfections

Cell transfections were done using Nucleofector II (Lonza, AAB-1001) with program 0-005 for hippocampal mouse neurons. The mouse neuron Nucleofector Kit (Lonza, VPG-1001) was used according to the manufacturer’s instructions. For each transfection, 5 * 10^5^ neurons were used with varying amounts for different plasmids: 15 or 25 μg of pBetaActin-TUBB2a-mEos3.2 and 15 or 25 μg of pBetaActin-TUBB2a-mEos3.2-mEos3.2-mEos3.2. All plasmids were purified using the EndoFree Maxiprep Kit (QIAGEN).

#### Live-cell imaging

All imaging experiments were performed on an Andor spinning disk Nikon Eclipse Ti microscope with Perfect Focus system (Nikon). The setup included a Yokogawa CSU-X1 spinning disk unit (CSUX1-A1N-E/FB2 5000 rpm Control, FW, DMB 95L100016) and a Plan Apo VC 60× oil objective NA 1.4 with no additional magnifier controlled via the iQ3 software (Andor). Single band-pass emission filters for the blue channel (480/40; Semrock), the green channel (525/50; Semrock BrightLine), and the red channel (617/73; Semrock BrightLine) were used. Excitation of the green channel was done with a 488-nm laser (100 mW), for the red channel with a 561-nm laser (100 mW) (REVOLUTION 500 series AOTF laser modulator and combiner unit, solid-state laser modules). Imaging was conducted inside a temperature-controlled large incubator chamber around the microscope, kept at 37°C. A small chamber for eight-well ibidi glass-bottom dishes (Pecon, #000470) was supplied with humidified air mixed with CO2 to maintain a concentration of 6–7%. Images were taken with an Andor iXON camera (DU897) in charge-coupled device mode at 1-MHz horizontal readout rate and with a pixel size of 0.22 μm and no binning.

The growth cone for measuring MT shrinkage speed (Extended data Fig. 2f-g) was imaged with a pixel size of 39 nm, time interval of 5 s and using a SORA super resolution spinning disk on a Nikon confocal spinning disc microscope.

Images were taken every 30 s for the first 4 min, followed by every 2 min as specified in the “Photoconversion” section.

For all experiments, neurons were selected based on representative morphology of more than two neurites. Imaging was done using high light exposure for high fluorescence intensity, which was essential to assess MT stability by mEos.

#### Photoconversion

Photoconversion was performed using the fluorescence recovery after photobleaching and photoactivation (FRAPPA) illumination system (Andor, Revolution). The FRAPPA module directs the laser onto individual pixels for a defined dwell time, sequentially scanning through all pixels for a predetermined number of repetitions. A predefined 4-pixel wide area was selected for photoconversion in a minor neurite (DIV1) and the axon (DIV3) to minimize phototoxicity while achieving high conversion intensities. Two photoconversions were done in the same minor neurite and axon. One was close to the soma and one far away from the soma, on average at 30% and 68% of the neurite length and 12% and 35% of the axon length. The patches were converted consecutively (approx. 2 h apart), ensuring that no significant signal from the previous conversion was visible anymore for a similar baseline before each conversion. This approach ensured that both conversions did not influence each others intensity measurements.

Photoconversion was performed with the 405-nm laser (800 repetitions, 20 μs dwell time, 1.78 μW power, pixel size of 0.22 μm, 10-pixel × 4-pixel). After photoconversion, images were acquired in the photoconverted (red) and not-photoconverted (green) channel every 30 s for the first 4.5 min, followed by every 2 min until most of the photoconverted mEos were depleted but at least 26 min in total. Exposure times were set at 800 ms at 90% or 100% laser intensity to achieve an optimal signal to noise ratio. The same imaging settings and filters were maintained across all experiments to ensure comparable fluorescence intensity levels. Most cells were re-evaluated 1 to 3 h after photoconversion to ascertain cell survival post-experiment; cells that died were excluded from the analysis.

#### Photoconversion analysis of experiments for MT decay curve

To measure photoconverted intensity, first the background value was determined. The background was calculated in the entire neuron, except in the neurite containing the photoconverted spot. In that area the 50^th^ highest pixel intensity value was measured for each timepoint and then the second highest value from all timepoints was increased by 2% and used as background value. Using this background value as threshold value to define the area with photoconverted intensity consistently worked well as judged by eye on most imaged neurons. This threshold value was subsequently subtracted from all intensity values and then the sum of all pixel intensity values in the neurite with the photoconverted spot was calculated for each timepoint. Once a noticeable part of the converted spot moved in the soma by eye, the analysis was stopped. This prevented photoconverted mass in the soma from affecting the analysis.

#### Photoconversion analysis of simulations for MT decay curve

To simulate the photoconversion experiment, for each MT the nucleation time, its x position, unstable length and stable length was saved. As before, simulations were first run until the steady state was reached and afterwards simulation data were saved each minute of simulation. After the simulation was fully done, the region where the photoconversion should be simulated was defined. At the first timepoint it was recorded for each MT how much length is within the defined range, and all lengths were summed to obtain the starting MT mass. Then for each timepoint for every MT that was lost (identified by the time it was nucleated), the corresponding length in the range was subtracted from the starting MT mass. As for photoconversion experiments, new growth of MTs or newly nucleated MTs were not considered. This allowed obtaining a curve of the remaining MT mass over time and determining the MT decay curve.

#### MT decay curve analysis

Data were fitted with the dual exponential function:

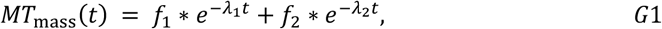

and for Fig. 1d also with the mono exponential function:

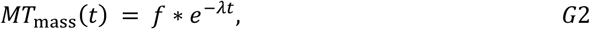

using scipy.stats.curve. For the dual exponential fit the fraction was calculated, e.g. for the fraction of population 1:

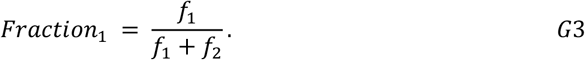

Cells for which the two fitted rates *λ*_1_ and *λ*_2_ were less than three-fold different, only one population of MTs was considered to be present. In general, the faster and slower decay rate were assumed to represent unstable and stable MTs, respectively. However, if the decay rate was found to be smaller than 0.23 min^-1^ for developing neurites (corresponding to a half-life of ca. 3 min) or smaller than 0.1 min^-1^ for dendrites and axons, the corresponding population was assumed to indicate stable MTs. If the decay rate was larger than these threshold values, the population was assumed to correspond to unstable MTs. For cases with only stable or only unstable populations present, an average decay rate was calculated:

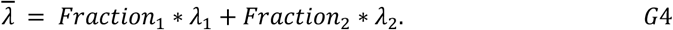

#### Measuring MT shrinkage speed

MT shrinkage speed was measured from single shrinkage events in one growth cone transfected with TUBB2a-mNeonGreen to be 10 µm/min ± 1 µm/min (n=5 events, N=1 growth cone; Extended data Fig. 2f-g). To account for MTs in the growth cone being pulled retrogradely due to actin retrograde flow, we reduced MT shrinkage speed by 3 µm/min, based on the measurement in^34^.

#### Immunocytochemistry

Neurons were incubated for either 24 or 72 h at 36.5°C and 5% CO_2_ and then fixed under conditions optimal for the preservation of the cytoskeleton: 4% paraformaldehyde, 4% sucrose in PHEM fixation buffer (60 mM Pipes, 25 mM HEPES, 10 mM EGTA, 2 mM MgCl_2_, pH 7.4) containing 0.1% Triton X-100 and 0.25% glutaraldehyde for 15 min. The coverslips with the cells were washed two times with PHEM buffer without fixatives and once with PBS. Fixed cells were quenched with 0.1 M glycine for 10 min, washed three times with PBS and then blocked at room temperature (RT) for 1 h with 2% Fetal Bovine Serum (FBS) (Thermo Fisher Scientific, A5256801), 2% Bovine Serum Albumin (BSA) (Sigma-Aldrich, A3294) and 0.2% fish gelatin (Sigma-Aldrich, G7765) in PBS. After blocking, cells were incubated with the primary antibodies (Tyrosinated Tubulin 1:800, Abcam ab6160; Detyrosinated Tubulin 1:500, Millipore MAB3201; Acetylated Tubulin 1:4000, Sigma T7451; Tubulin, alpha, (clone B-5-1-2) mouse monoclonal Ab, Sigma T5168, 1:4.000) for 1 h at RT. Cells were washed four times with PBS and incubated with the secondary antibodies (1:1000) (Alexa-Fluor 488 *α*-Rat, Invitrogen, A21208; Alexa-Fluor 555 *α*-Rabbit, Invitrogen, A21429; Alexa-Fluor 555 *α*-Mouse, Invitrogen, A21422; Alexa-Fluor-488 anti-Mouse, Invitrogen, A11029), for 30 min at RT. The antibodies were mixed in a 1:10 dilution of the blocking solution in PBS. Cells were washed three times with PBS, once with ddH_2_O and mounted onto microscope slides with Fluoromount^™^ (Sigma, F4680).

#### Imaging tubulin modifications in fixed cells

Cells for imaging of MT distribution were selected randomly. Cells were imaged using an Axio Observer.Z1 microscope equipped with a Colibri 7 LED light source and Zen Blue software (all from Zeiss). Images were taken with an Axiocam 512 mono camera using either a Plan Apochromat 63x oil objective numerical aperture (NA) 1.4 with a pixel size of 0.098 µm (2 x 2 binning) or a Plan-Apochromat 40x oil objective NA 1.4 with a pixel size of 0.078 µm (no binning). A dichroic Quad filter (QBS 405, 493, 575 and 653) and single band-pass emission filter for the green channel (525/50) and the red channel (605/40) were used. For the green and red channel, a 475 nm LED-module and a 555 nm LED-module were used.

#### Experimental MT distribution analysis

To analyze the experimental MT distribution, neurites were traced in FIJI. The traces were used in python to measure the summed intensity across the width of the neurite (up to a maximum of 31 pixels for the shaft and 71 pixels for growth cone) along the traced line. Images were background subtracted, with the background value identified for each cell separately as the highest value outside but close to the cell. For this a manual region close to the cell was drawn in FIJI and read out in python.

To compare MT decay data with experimental MT distributions (Fig. 1q and Extended Data Figs. 1e, 7i), a normalized ratio change, 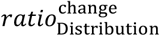, was calculated for both MT distributions and MT decay. This ratio change is invariant to the absolute ratio of stable to unstable MTs:

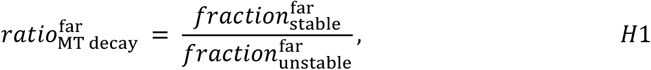

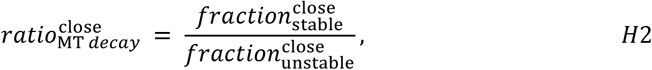

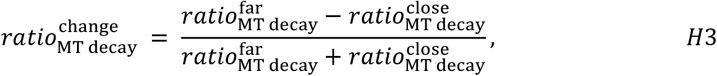

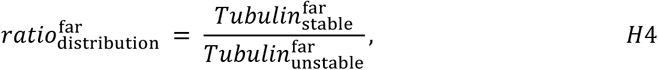

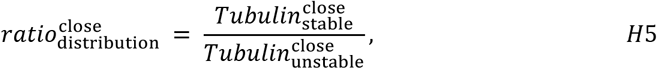

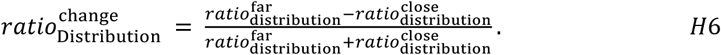

#### Calculating possible distributions for non-acetylated tubulin

Modifications of tubulin give an indication of the age of a MT – with acetylation and detyrosination accumulating on older MTs. Tyrosinated MTs should indicate young MTs and thus the distribution of tyrosinated MTs should be different from the distribution of acetylated and detyrosinated MTs. Instead, we found that the distribution for acetylated tubulin looked more similar to tyrosinated tubulin than to detyrosinated tubulin (Extended Data Fig. 1b). This indicates that the distribution of acetylated MTs contains a higher fraction of tyrosinated MTs than of detyrosinated MTs. Thus, the distribution of tyrosinated MTs also contains a substantial fraction of older MTs (stable MTs) that are acetylated. If the amount of MT modifications depends on the age of MTs, this would indicate that acetylation occurs noticeably faster than detyrosination – since otherwise tyrosinated MTs would not be acetylated often. Indeed, in non-neuronal cells, MTs accumulate a lot of acetylation within 15 min after MT regrowth, but it takes more than 90 min to accumulate considerable fractions of detyrosination^26^. This suggests that most MTs that are non-acetylated are tyrosinated and not detyrosinated, since they are not old enough to accumulate detyrosination. In summary, the distribution of non-acetylated should label MTs that are on average younger than tyrosinated MTs, since even some older MTs that are acetylated should still be tyrosinated. Thereby, non-acetylated tubulin could be a better marker for young, unstable MTs. To calculate possible distributions for non-acetylated MTs (that are also tyrosinated), average distributions for detyrosinated, tyrosinated and acetylated tubulin were used. Since acetylation/non-acetylation and detyrosination/tyrosination can occur together but MTs cannot be acetylated and non-acetylated or detyrosinated and tyrosinated at the same time, the following four combinations of modifications are possible for MTs: acetylated and tyrosinated, acetylated and detyrosinated, non-acetylated and tyrosinated as well as non-acetylated and detyrosinated. We measured the following three distributions experimentally:

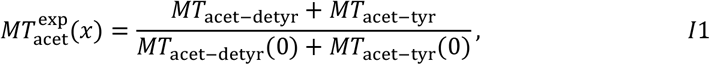

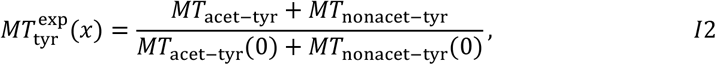

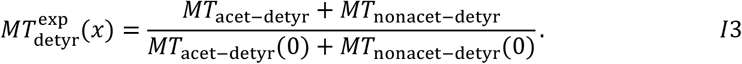

Since the fluorescent signals from the three markers are arbitrary units, we normalized each distribution by the value in the neurite close to the soma (x=0). Therefore, the experimental measurements only allow us to obtain a relative distribution and leaves the ratio of the three distributions unknown – e.g. we cannot measure how much more acetylation there is compared to detyrosination.

Since detyrosination happens a lot slower than acetylation in dividing cells^26^, we don’t expect MTs that are detyrosinated to still be nonacetylated. Thus, we assumed that the ratio of MTs that are non-acetylated and detyrosinated to MTs that are acetylated and detyrosinated is close to 0:

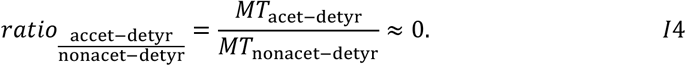

Thus, the measured distribution of detyrosinated MTs should represent the distribution of detyrosinated MTs that are also acetylated:

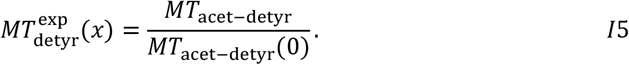

We now assume different fractions of acetylated MTs that are also detyrosinated (0.05, 0.1 and 0.15) in the neurite close to the soma (x=0):

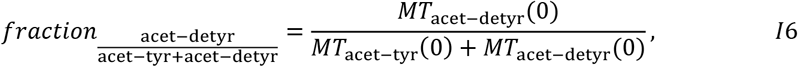

and different fractions of tyrosinated MTs that are also acetylated (5%, 15%, 25%, 35%, 45%, 55%, 65% and 75%) in the neurite close to the soma (x=0):

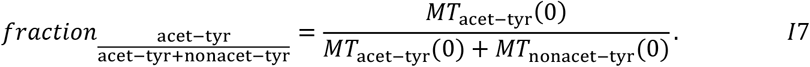

We combine equations (I5), (I7) and (I1) to use the 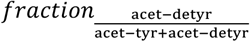 for calculating the distribution of MTs that are acetylated and tyrosinated:

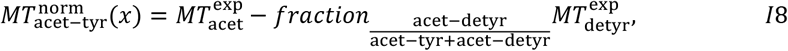

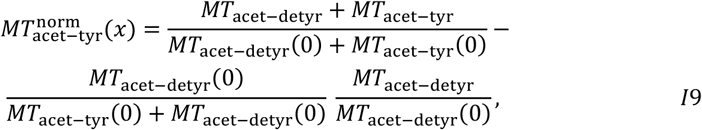

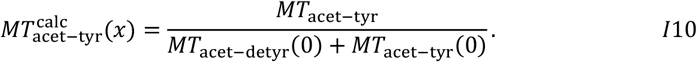

We normalize equation (I10) by its value in the neurite close to the soma, to obtain the normalized distribution of MTs that are acetylated and tyrosinated:

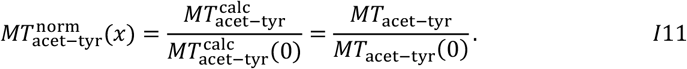

Finally, we combine equations (I2), (I7) and (I10) to subtract this distribution from the distribution of tyrosinated MTs, resulting in normalized distribution of non-acetylated MTs that are tyrosinated, which is the distribution of non-acetylated MTs we used as marker for unstable MTs:

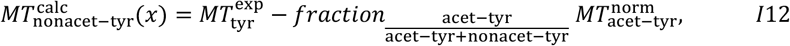

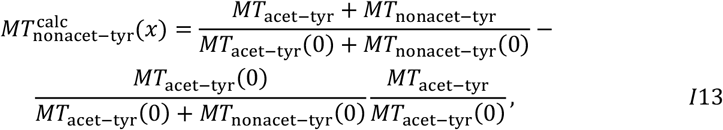

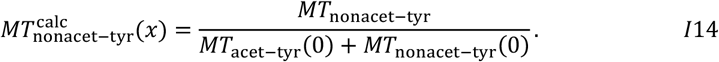

Lastly, by normalizing the distribution by its value in the neurite close to the soma (x=0), we obtain the normalized distribution of nonacetylated MTs that are tyrosinated:

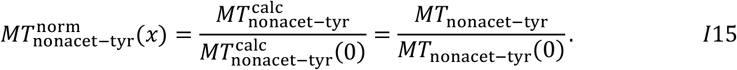

#### Measuring the number of MTs at the neurite base

Estimating the number of MTs at the neurite base was done similarly as in^34^. The measured total intensity of alpha Tubulin was divided by the intensity of a single MT, to obtain an estimate for the number of MTs (Extended Data Fig. 2d,e). Single MTs could not be identified visually in the neurite or soma, but in the growth cone. For these single MTs, the intensity of one pixel length was measured across the entire MT. All intensity values for one pixel length MT from one cell were averaged. The intensity of alpha Tubulin at the neurite base was then normalized by this average single MT intensity per pixel length.

#### Analysis of EB3 lifetime and distribution

Live-cell imaging movies of EB3 were processed automatically with the “plusTipTracker” of “u-track 2.0” implemented in Matlab^77^. For detecting comets, a “low-Gaussian SD” of 1 pixel, “high-pass Gaussian SD” of 4 pixels, and a “minimum threshold” of 3 SDs with a “threshold step size” of 1 SD were employed. To track comets, these parameters were used: “maximum gap to close” was 2 frames, “minimum length of TrackSegments from First Step” was 3 frames, with segments being split. For “Cost functions” and “Kalman filter functions” the option “Microtubule plus-end dynamics” was used. To obtain the distribution of EB3 comets, further analysis of comet tracks was performed using the previously developed Python package “neuriteMT”^16^ (https://github.com/maxschelski/neuritemt). This package builds an image of the neuron using all comet tracks identified from all timepoints. This neuron image was then processed further using the pyNeurite package^16^, which created sorted skeleton points of all neurites. For each timepoint a distribution of the number of EB3 comets per µm was obtained for all neurites. EB3 comets attributed to the first pixel of the neurite were excluded to remove EB3 comets from the soma. These distributions were averaged across all neurites from 20 µm to 40 µm long to obtain the average distribution of EB3 comets in the neurite.

To obtain the lifetime of EB3 comets (Extended data Fig. 2a-c), we calculated the histogram of the measured EB3 lifetimes (seconds they were present) and fitted a mono-exponential function to it using the curve_fit function in the python package scipy.optimize:

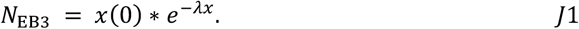

Where *N*_*EB*3_ is the number of EB3 comets with a specific lifetime (in the histogram bin) and x is the measured lifetime of EB3 comets. The fitted lifetime 1/*λ* was used for the analysis. Histograms were obtained for each neuron and the average of the lifetimes of all cells was used.

#### Analysis of MT loss for MT-RF speedup experiments

First, the distribution of MTs was analyzed, as defined in “Experimental MT distribution analysis”. Then either the average intensity in the entire neurite (global) or the intensity from 60% to 90% neurite length towards the neurite tip (distal) was analyzed for each timepoint.

For neurites, tracing was done once for all timepoints, to analyze local MT mass in the neurite for each timepoint. The first timepoint after the treatment started (clearly before MT-RF speedup) was chosen as the starting value. The second timepoint was separately chosen for global and distal MT mass as the lowest value within 15 min after treatment started. Timepoints for the analysis of simulations were chosen the same way.

For dendrites, due to massive changes in neurite shape, local dendritic MT mass was only analyzed at two timepoints, by tracing dendrites separately at these two timepoints. The baseline timepoint was the first timepoint after the treatment had started, as for neurites. To measure MT loss, the second timepoint was chosen by eye to be within 15 min after the timepoint with the fastest visible MT-RF speed. Due to variable onset times of visible MT-RF speedup and variable MT-RF acceleration, the corresponding timepoints varied between 12 and 144 min. For simulation data, to match the experimental timeframe, the timepoint with the lowest number of MTs either globally or distally within 60 min after MT-RF speedup was chosen. This way, the average timeframe used for the analysis of simulation data for the entire neurite (global) was 51 min after speedup, exactly like for experimental data.

#### Python scripts

All scripts used in the manuscript were written with Python 3.7 or 3.8. Environments were constructed with conda or mamba using the conda-forge channel (Anaconda Inc., 2020; available from https://docs.anaconda.com/; https://github.com/mamba-org/mamba). The following packages were used: SciPy 1.6.2 (for FigureFlow) or 1.5.2 (https://pypi.org/project/scipy/)^78^, sci-kit image 0.17.2 or 0.18.1 (https://pypi.org/project/scikit-image/)^79^, numpy 1.20.3 (https://pypi.org/project/numpy/)^80^, seaborn 0.11.1 (for FigureFlow) or 0.11.0 (https://pypi.org/project/seaborn/)^81^, scikit-posthocs 0.6.7 (https://pypi.org/project/scikit-posthocs/)^82^, matplotlib 3.34 or 3.4.2 (https://pypi.org/project/matplotlib/)^83^, pandas 1.3.0 (for FigureFlow) or 1.1.3 (https://pandas.pydata.org/)^84^, python-pptx 0.6.19 (https://pypi.org/project/python-pptx/). For cytostoch the following packages were used: numba 0.60 (https://github.com/numba/numba), pytorch 2.2.2 (https://pytorch.org/), pandas 2.1.1 (https://pandas.pydata.org/)^84^, pyarrow 11.0 (https://pypi.org/project/pyarrow/), matplotib 3.8.0 (https://pypi.org/project/matplotlib/)^83^, seaborn 0.12.2 (https://pypi.org/project/seaborn/)^81^, dill 0.3.7 (https://pypi.org/project/dill/).

#### Figure preparation (FigureFlow)

All figures were prepared using our Python package FigureFlow^16^ . The package is available via GitHub (https://github.com/maxschelski/figureflow).

#### Statistics

Generally, only statistical comparisons between experimental data are shown. We decided not to perform statistical comparisons between simulation data and experimental data for two reasons. First, we do not want to claim that experimental data and simulation data are identical. Instead, we want to highlight that our simulations often capture the main features of experimental data and usually are quantitatively close to experimental data. Second, the large number of simulations for model data can make even very small differences significant. This can lead to significant differences, when distributions by eye seem to not differ a lot. Instead of statistical comparisons, for all data, except distributions across the neurite, we show boxplots of all simulation data and experimental data to allow a good comparison between model and experimental data by eye. This allows to judge not only how big the difference of the averages is but also how much of the variance in experimental data is captured by the stochastic variations in the model. For all data where more than two groups were compared, the nonparametric Kruskal-Wallis test (scipy.stats.kruskal_wallis) was first performed. If the resultant *P* value was below 0.05, all combinations of groups were compared. All statistical comparisons of the two groups were undertaken using the nonparametric Dunn’s test (scikit-posthocs.posthoc_dunn), with Bonferroni correction. If only two groups were compared, the Mann-Whitney U test was used.

## Data availability

Previously acquired experimental data that was analyzed in this study is publicly available (http://doi.org/10.5061/dryad.2fqz612s8). Experimental data newly acquired for this study and model data will be made publicly available.

## Code availability

The Python package to run stochastic simulations of MTs in neurites developed for this study is available from Github (https://github.com/maxschelski/cytostoch). Further Python packages used in this study are publicly deposited (http://doi.org/10.5281/zenodo.6990848). All scripts used in this study to generate figures and data will be publicly deposited.

## Acknowledgements

We thank J. Schiweck, A. Husch, S. A. Fiorenza, C. Bergmann and I. Tolic for critically reading and discussing the manuscript; and the Animal Research Facility, the Light Microscope Facility of the DZNE Bonn, A.-T. Pham, B. Randel and J. Benner for technical assistance. This work was supported by the Deutsche Forschungsgemeinschaft (DFG), the International Foundation for Research in Paraplegia (IRP), and Wings for Life. M.S. was supported by the Add-On Fellowship for Interdisciplinary Sciences from the Joachim Herz Foundation and a fellowship from the International Max Planck Research School for Brain and Behavior (IMPRS-BB). C.G. is a recipient of the Drs. Ayeez and Shelena Lalji & Family Student Scholar Award for Repair and Regenerative Mechanisms in ALS. F.B. is a member of the excellence cluster ImmunoSensation2, the SFBs 1089, 1158 and 1690, the SPP 2395, and the NRW network iBehave, funded by the Chan-Zuckerberg Initiative (CZI) and is a recipient of the Roger De Spoelberch Prize and the Drs. Ayeez and Shelena Lalji & Family Award for Innovative Healing.

## Author Information

### Contributions

M.S. conceived the project. M.S., N.P. and F.B. designed the research. M.S. and N.P. developed the theory. M.S., T.P., C.G. and S.S. performed the experiments. M.S. and C.G. performed analysis of raw images. M.S. analyzed the data. M.S. developed the software. M.S. N.P. and F.B. supervised the research. M.S., N.P. and F.B. wrote the manuscript. C.G., T.P. and S.S. reviewed the manuscript.

---

**Extended Data Fig. 1.**
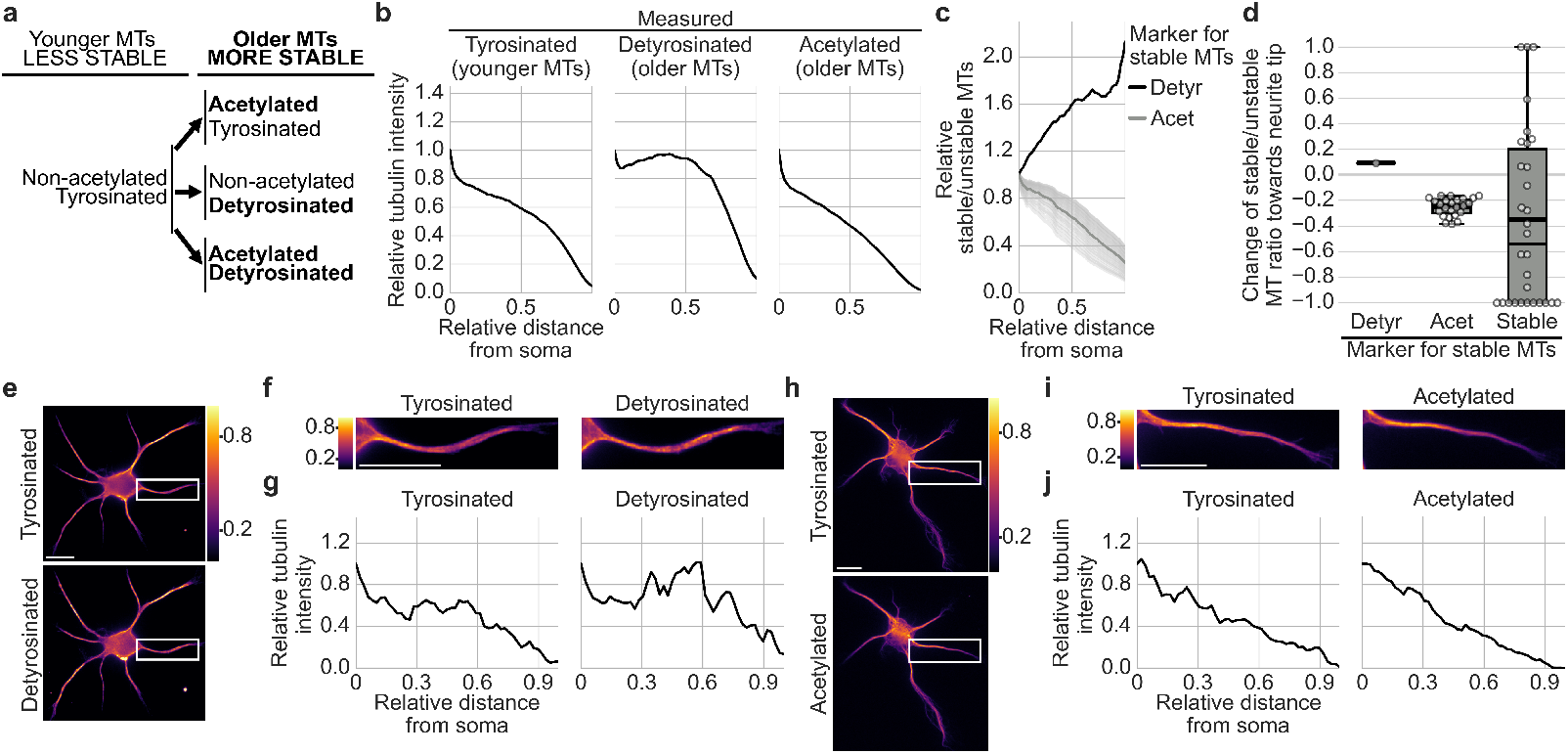
Acetylation is an appropriate marker for stable microtubules. Neurons were stained after one day in culture for tyrosination as a marker for younger and detyrosination as well as acetylation as a marker for older MTs. Intensities were quantified for each point along the neurite as the sum of all pixel intensities across the width of the neurite. Tubulin intensity was normalized to the value closest to the soma. (For Tyrosinated: n = 1144 neurites from 331 neurons, N = 6; Detyrosinated: n = 639 neurites from 188 neurons, N = 3; Acetylated: n = 505 neurites from 143 neurons, N = 3 experiments). **a**, Illustration of MT modifications that neurons were stained for. **b**, Distributions of tubulin modifications were averaged for all neurites. **c**, The distributions of the relative ratio of stable to unstable MTs for detyrosinated tubulin (Detyr) or acetylated tubulin (Acet) as marker for stable MTs. **d**, The normalized change of the ratio of stable to unstable MTs from close to the soma to far from the soma was calculated and normalized by the sum of the ratios at both positions. Detyrosinated and tyrosinated (Detyr) or acetylated and non-acetylated (Acet) tubulin were used as markers for stable and unstable MTs, respectively. This was compared to the value obtained from the data in Fig. 1p as a good marker for stable MTs (Stable). **e-j**, Representative images (**e,h**) of the entire neurons and (**f,i**) the region marked with the white rectangle enlarged, and (**g,j**) the intensity curve along the neurites from **f,i**, stained for (**e,f**) tyrosinated and detyrosinated or (**i,j**) tyrosinated and acetylated tubulin. Scale bars, 10 μm. Intensity in **e,f,h,i** is color-coded from purple (low) to yellow (high). The thick line in boxplots indicates the mean, the thin line the median.

**Extended Data Fig. 2.**
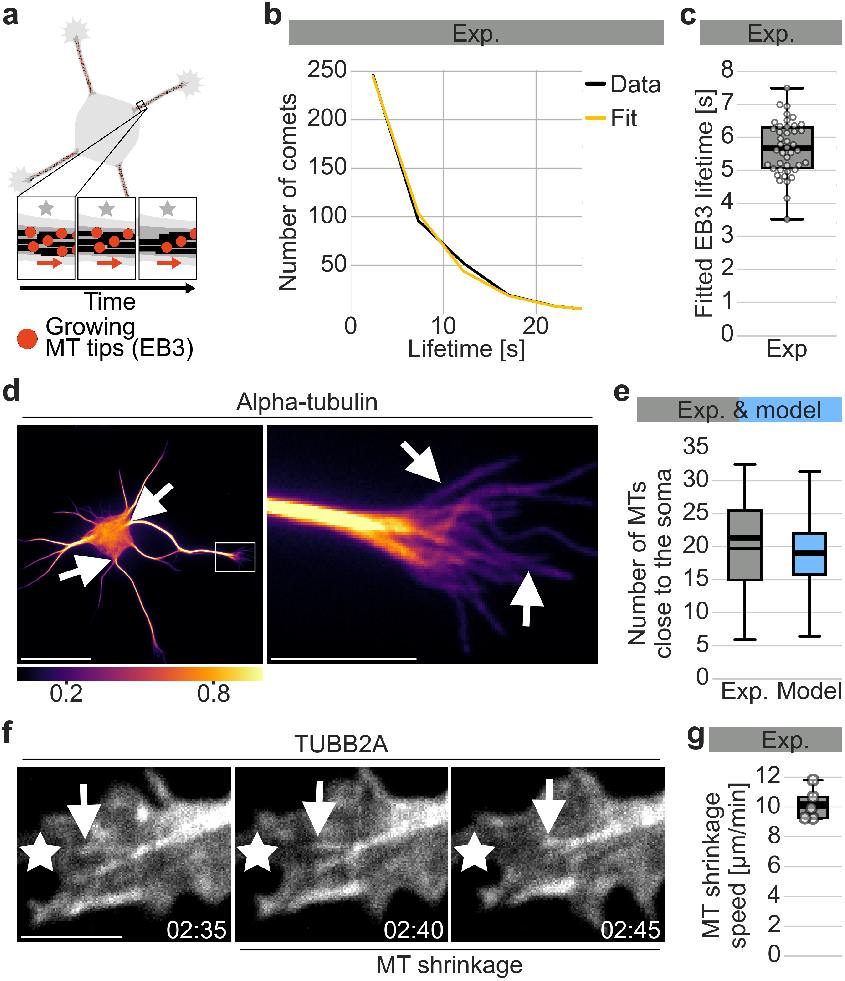
Measuring the pausing rate, neurite base MTs and retraction speed. **a-c**, Neurons expressing fluorescently labeled EB3 were expressed and imaged after one day in culture in ^16^. (**a**) Illustration of imaging growing MT tips. (**b,c**) The histogram of EB3 comet lifetimes was measured from data from ^16^. The histogram (Data) was fitted with a mono-exponential function (Fit). (**b**) Histogram of EB3 lifetime for a representative neuron in **c**. (**c**) Fitted EB3 lifetimes (n = 42 neurons, N = 3 experiments). **d,e**, Neurons grown for one day in culture were fixed, stained for alpha-tubulin and imaged. The average intensity of single MTs in growth cones per pixel was measured. The intensity of the neurite at the soma along a one-pixel-wide line was summed and normalized by the average intensity of a single MT per pixel to obtain an estimate of the number of MTs close to the soma. (**d**) Representative neuron with the neurite close to the soma indicated by white arrows on the left side and single MTs in the growth cone indicated by white arrows in the right zoom. Intensity is color-coded from purple (low) to yellow (high). Scale bars, 20 µm for not zoomed and 5 µm for zoomed image. (**e**) Comparison of experimentally measured number of MTs close to the soma and the number of MTs in the model. Simulation data from Fig. 3g (Exp: n = 32 neurons, N = 1 experiment; model: n = 16000 simulations). Due to the large number of simulations, model and experimental data are not statistically compared (Methods section “Statistics”). **f,g**, The growth cone of a neuron expressing the fluorescently labeled tubulin subtype TUBB2a was imaged after one day in culture. (**f**) White arrows indicate a shrinking MT and the star a fixed position. Time is in min:seconds. Scale bar, 2 µm. (**g**) MT shrinkage events were analyzed from the imaged growth cone (n=5 events, N=1 experiment). The thick line in boxplots indicates the mean, the thin line the median.

**Extended Data Fig. 3.**
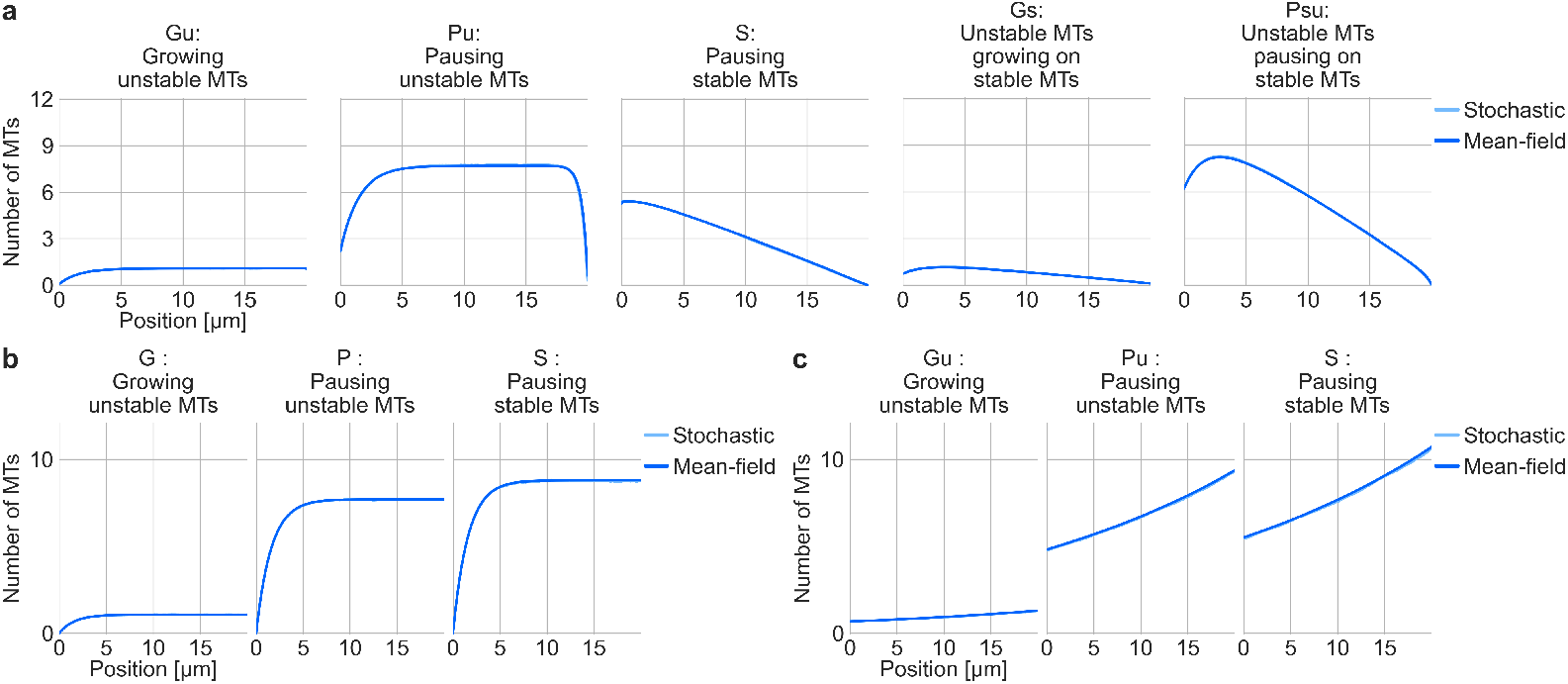
Stochastic simulations match mean-field solutions. Mean-field steady-state solutions were calculated from systems of partial differential equations. The equivalent stochastic simulations were run until steady state (n = 4000 simulations). **a**, Somatic and MT branching nucleation rates were set to zero. Mean-field solutions were obtained numerically. **b,c**, MT-RF speed was set to zero. Additionally, stable growth rate was set to zero to prevent the formation of unstable MTs growing or pausing on stable MTs. (**b**) Somatic and MT branching nucleation rates. Mean-field solutions were obtained analytically. (**c**) Somatic nucleation rate was set to zero, but MT branching rate was nonzero and not limited by resource availability. Mean-field solutions were obtained numerically.

**Extended Data Fig. 4.**
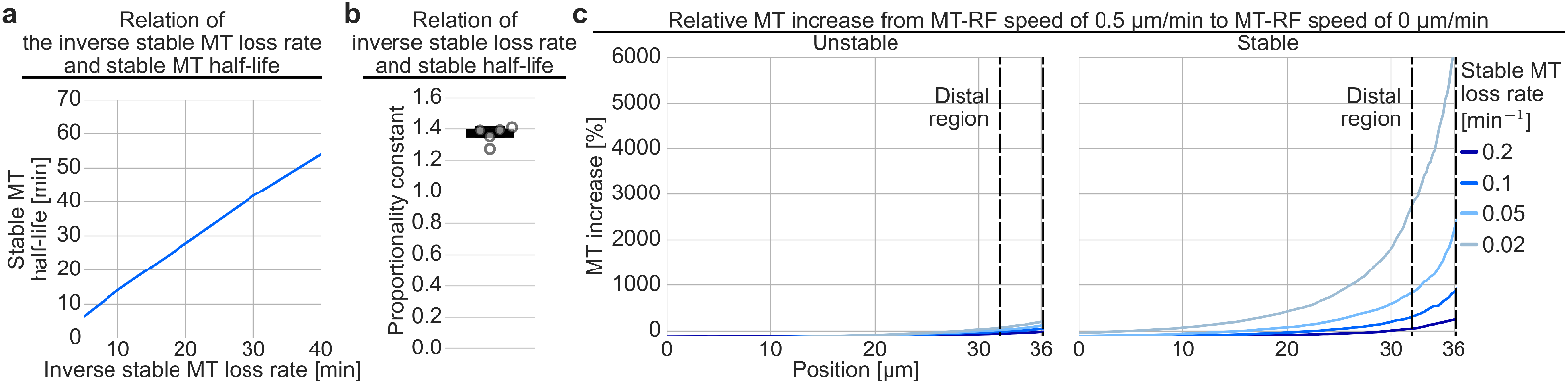
Stable MT half-life correlates with inverse stable loss rate and MT-RF slowdown increases MTs the most distally. Simulations over 40 µm with parameters as in Figure 2. **a**, The fitted stable MT half-life is shown for simulations run with different values for the inverse stable MT loss rate. **b**, The proportionality constant of the fitted stable MT half-life divided by the inverse of the stable MT loss rate is shown. (n = 2000 simulations, simulation time: 510 min). **c**, The increase in unstable and stable MTs without MT-RF compared to 0.5 µm/min MT-RF speed is calculated from data in Fig. 2c, d (n = 2000 simulations, simulation time: 450 min).

**Extended Data Fig. 5.**
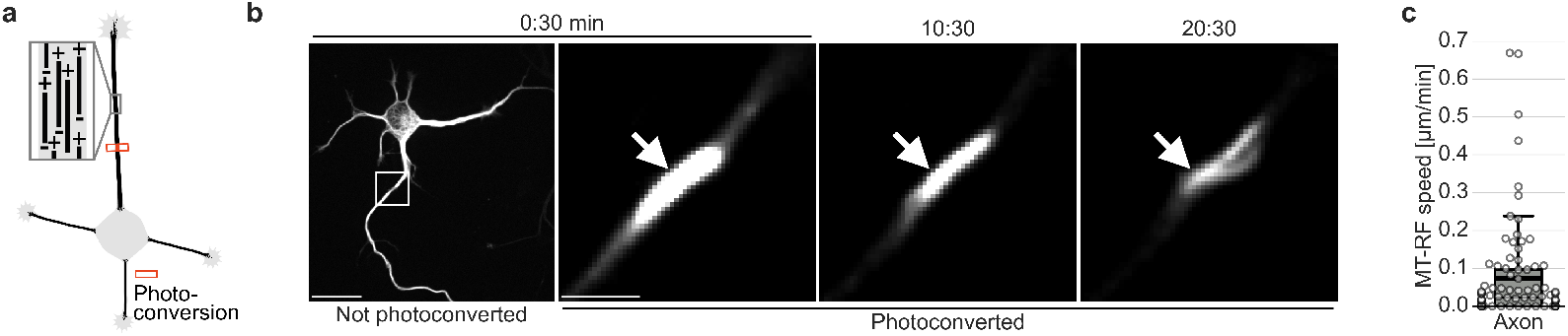
MT-RF slows further down in old axons. Neurons expressed the tubulin TUBB2a fused to the photoconvertible fluorophore mEos3.2 and were cultured for three days. Then a small MT patch in the axon was photoconverted on average at 32% neurite length towards the axon tip. **a**, Illustration of photoconversion experiments. **b**, Representative neuron for **c**. The white arrow points to the position of photoconversion. Time is in min:seconds. Scale bars, 20µm for not zoomed and 4µm for zoomed image. **c**, MT-RF was measured (n = 46 neurons, N = 4 experiments). The thick line in boxplots indicates the mean, the thin line the median.

**Extended Data Fig. 6.**
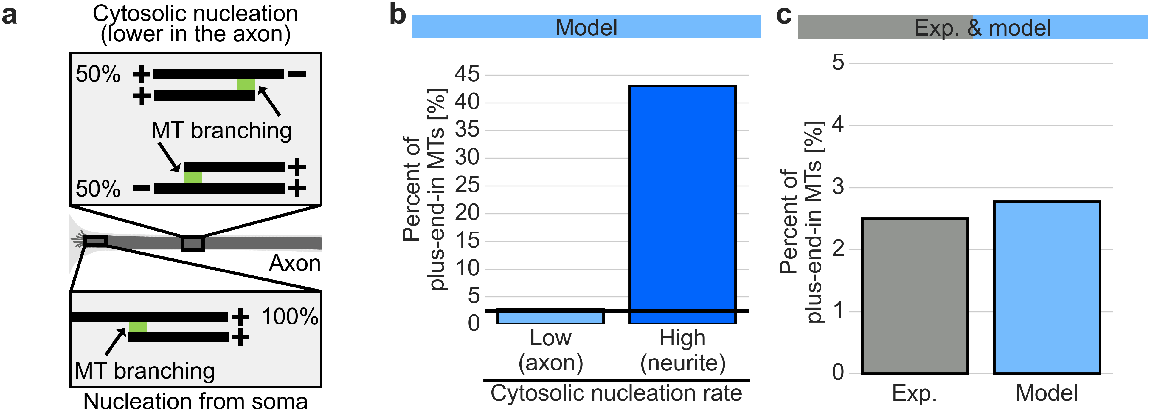
Percent of model plus-end-in MTs with varying cytosolic nucleation rates. Simulations with low (0.003 min^-1^µm^-1^) or high (0.2 min^-1^µm^-1^) cytosolic nucleation rate with slow MT-RF of 0.1 µm/min are run (n = 500 simulations, simulation time: 1600 min). **a**, Illustration of the influence of nucleation mechanisms on MT orientation and the reduction of cytosolic MT nucleation used in simulations of axons. **b**, The theoretical percentage of plus-end-in MTs is analyzed. The horizontal black line indicates the experimentally expected value of 2.5%. **c**, Theoretical percentage of plus-end-in MTs in simulations with reduced cytosolic nucleation rate (from 0.2 min^-1^µm^-1^ to 0.003 min^-1^µm^-1^) compared to experimental data of 2.5% plus-end-in MTs from^5^ (n = 500 simulations, simulation time: 1600 min).

**Extended Data Fig. 7.**
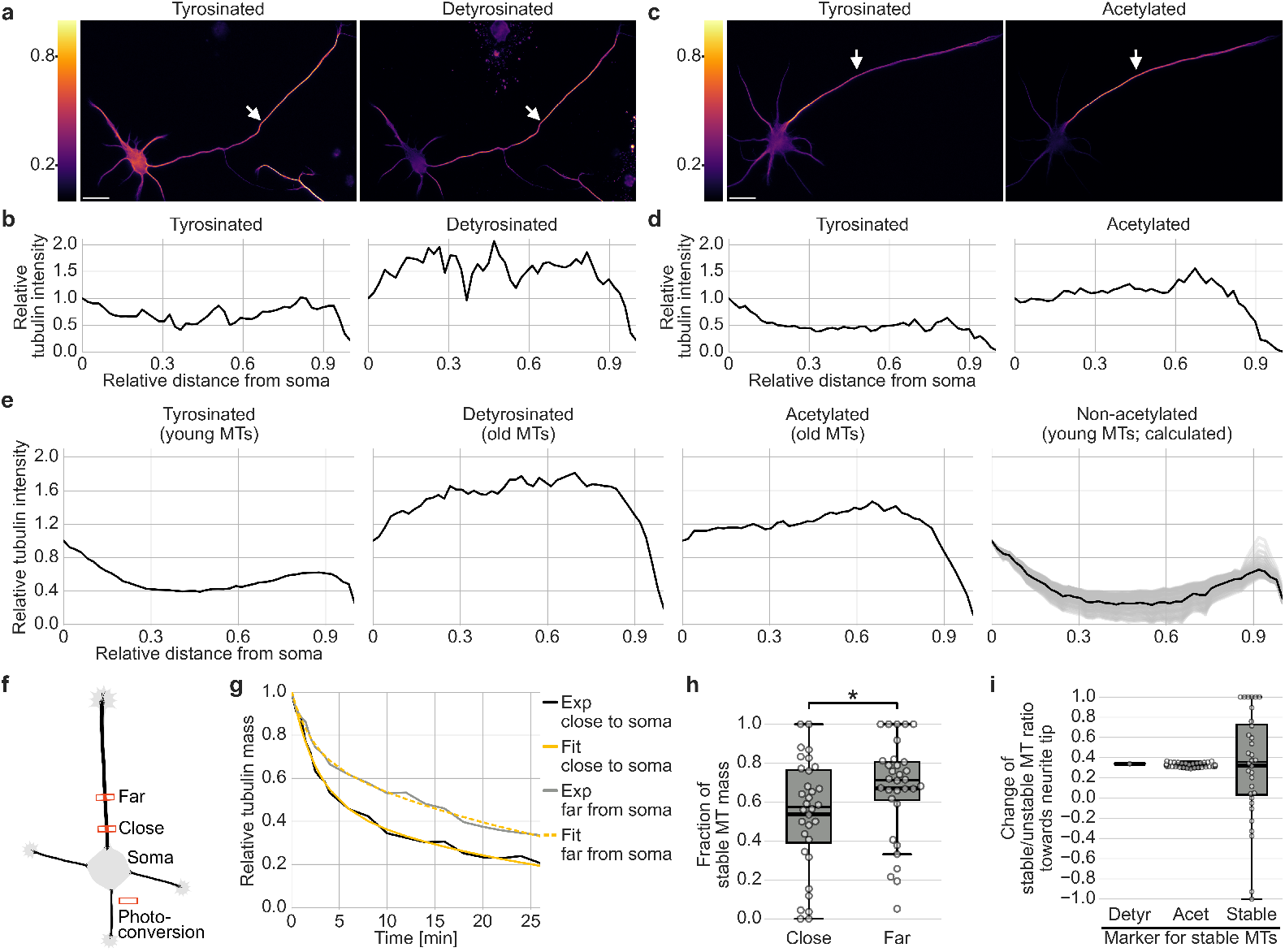
Distributions of stable and unstable MTs in axons. **a-e**, Neurons cultured for three days were stained for (**a,b**) tyrosinated and detyrosinated or (**c,d**) tyrosinated and acetylated tubulin and the corresponding distributions were measured for 100 µm to 200 µm long axons. Intensities were quantified for each point along the axon by summing up all pixel intensities across the axon width at that point and normalizing the value by the point closest to the soma. Intensity in **a,c** is color-coded from purple (low) to yellow (high). (**a-d**) (**a,c**) Images and (**b,d**) intensity curves of representative neurons for **e**. Scale bars, 20 µm. (**e**) Distributions were averaged across axons of all neurons by normalizing the position along the axon. The non-acetylated distributions were calculated from the other distribution as in Fig. 1l (Tyrosinated: n = 114 neurons, N = 4 experiments; Detyrosinated: n = 58 neurons, N = 4 experiments; Acetylated: n = 56 neurons; N = 3 experiments). **f-h**, TUBB2a-mEos3.2 expressing neurons were cultivated for three days, after which an MT patch close to or far from the soma in the axon was photoconverted, on average 12% and 35% axon length towards the axon tip, respectively. For each axon a close and far MT patch were converted separately. (**f**) Illustration of photoconversion experiment. (**g,h**) The photoconverted intensity was measured for each timepoint and a dual exponential function fitted to the decay curve. (**g**) Decay curves for representative neuron. (**h**) Fraction of stable MT mass close to and far from soma (n = 32 neurons, N = 4; p = 0.021, Mann-Whitney U test). **i**, Comparison of the normalized change of the stable/unstable MT ratio from close to the soma to far from the soma for photoconversion experiment (Stable) and when using detyrosinated and tyrosinated tubulin (Detyr) or acetylated and non-acetylated tubulin (Acet) as markers for stable and unstable MTs, respectively. The thick line in boxplots indicates the mean, the thin line the median.

